# Convergence of dispersed regulatory mutations predicts driver genes in prostate cancer

**DOI:** 10.1101/097451

**Authors:** Richard C. Sallari, Nicholas A. Sinnott-Armstrong, Juliet D. French, Ken J. Kron, Jason Ho, Jason H. Moore, Vuk Stambolic, Stacey L. Edwards, Mathieu Lupien, Manolis Kellis

## Abstract

Cancer sequencing predicts driver genes using recurrent protein-altering mutations, but detecting recurrence for non-coding mutations remains unsolved. Here, we present a convergence framework for recurrence analysis of non-coding mutations using three-dimensional co-localization of epigenomically-identified regions. We define the regulatory plexus of each gene as its cell-type-specific three-dimensional gene-regulatory neighborhood, inferred using Hi-C chromosomal interactions and chromatin state annotations. Using 16 matched tumor-normal prostate transcriptomes, we predict tumor-upregulated genes, and find enriched plexus mutations in distal regulatory regions normally repressed in prostate, suggesting out-of-context de-repression. Using 55 matched tumor-normal prostate genomes, we predict 15 driver genes by convergence of dispersed, low-frequency mutations into high-frequency dysregulation events along prostate-specific plexi, while controlling for mutational heterogeneity across regions, chromatin states, and patients. These putative drivers play roles in growth signaling, immune evasion, mitochondrial function, and vascularization, suggesting higher-order pathway-level convergence. We experimentally validate the *PLCB4* plexus and its ability to affect the canonical PI3K cancer pathway.

## INTRODUCTION

Sequencing has revealed both germline variants that underlie cancer risk and somatic mutations that drive cancer progression. However, disparate perspectives emerge from each. Germline variants identified in genome-wide association studies (GWAS) are predominantly distal to genes. When experimentally characterized they appear to interact with genes by folding to their promoter’s physical location in three-dimensional space. These variants have subtle effects on transcription factor binding and gene expression but ultimately stack the odds towards disease development or progression. By contrast, somatic alterations identified through tumor sequencing are far more severe and have a direct impact on protein sequence. Both emerging perspectives, the germline and the somatic, belong to the same disease, but it is not yet clear how they are related.

Here, we present a unifying framework to reconcile these disparate perspectives and venture into the continuum between them. In this framework we propose the existence of a theoretical genomic event in which heterogeneous variants that are scattered and far from each other on the one-dimensional genome sequence but that are physically adjacent to each other in the three-dimensional volume of the cell nucleus manifest as a coherent cellular phenotype. We define a plexus as a set of interacting loci that are next to each other in the cell volume but scattered over the genome sequence. The number of possible plexi quickly becomes astronomical, even when they are composed of a handful of loci. Without experimental knowledge of the location of active loci and their interactions, the plexus framework is computationally and statistically intractable. Minimally, looking for a driver plexus requires whole genome sequencing of cancer-normal pairs and maps of chromatin states and chromosome interactions for the cancer’s tissue of origin.

In this study, we apply the plexus framework to prostate adenocarcinoma. We use matched ChIP-seq and Hi-C to map the locations of regulatory elements and their interactions in normal prostate cells (RWPE1). We then build the prostate-specific plexus for every protein-coding gene in the human genome. We initially use the plexi to analyze dysregulated genes in 16 cancer-normal transcriptome pairs. This approach reveals that the plexi of dysregulated genes enrich in dispersed non-coding mutations that converge on gene promoters and disrupt their function. We then develop a plexus recurrence test that we apply to 55 cancer-normal whole genome pairs. This test allows us to uncover 15 driver plexi containing novel candidate cancer genes with diverse roles that further converge on growth signaling, immune evasion, mitochondrial function and vascularization. Finally, we experimentally demonstrate how our most robust result, the *PLCB4* plexus, disrupts the PI3K pathway to alter cell growth and drive tumor progression. Our results have broad implications beyond identifying cancer genes. We hope the plexus framework will boost power in sequence association studies and facilitate the interpretation of rare and private variants in the context of precision medicine.

## DESIGN

### The plexus framework

The leading paradigm for identifying driver genes in cancer genomics has been to search for recurrent mutations in multiple independent patients. This process is confounded by many factors, such as the variation in mutation rates across the genome due to transcription and replication timing (Lawrence et al. 2013). While cancer recurrence has typically focused on protein-coding mutations, recent studies show that regulatory regions can also be the targets of recurrent mutations, in particular, single regulatory regions linked to the *TERT* (Huang et al. 2013) and *TAL1* genes (Patel et al. 2014). But in contrast to coding recurrence, where hundreds of genes have been identified and many more remain to be discovered (Lawrence et al. 2014), the analysis of recurrence at single regulatory regions has not yielded conclusive results (Nik-Zainal et al. 2016).

Expanding the concept of recurrence to non-coding mutations poses several challenges that are currently unmet. First, cancer is a highly heterogeneous disease among patients. Because multiple regulatory loci can be associated with the same gene, each locus might be mutated in a different tumor sample, requiring methods that go beyond a single region. Second, regulatory loci can lie far from the genes they regulate. This demands precise methods for identifying interacting loci over long-range chromatin conformation loops (Pomerantz et al. 2009; Ahmadiyeh et al. 2010; Zhang et al. 2012). Third, mutation rates are heterogeneous over the genome. Regions associated with active histone modification marks, for example, can show dramatically lower background mutation rates due to higher accessibility for the DNA repair machinery (Stamatoyannopoulos et al. 2009; Liu, De, and Michor 2013; Polak et al. 2014). Fourth, the regulatory code is poorly understood. This complicates the prioritization of mutations and the assessment of regulatory consequences (Cowper-Sal·lari et al. 2012; Ernst and Kellis 2013; Khurana et al. 2013). Fifth, all the parameters used in the statistical modeling vary by cell type. The interactions between loci, chromatin states, DNA accessibility and the concentrations of the transcription factors decoding the regulatory instructions are specific to each and every cell type in the human body.

Here, we directly address these challenges and introduce a theoretical and methodological framework for recurrence analysis of non-coding mutations that we apply to prostate cancer. We begin by inferring the plexus of every protein-coding gene (Fig. 1a), defined as the set of all proximal and distal regulatory elements acting through intra- or inter-chromosomal interactions (also *cis* and *trans*, respectively) based on their chromatin state and the three-dimensional links to their target genes. This allows us to collapse mutations that are heterogeneous across samples and scattered over multiple genomic loci (Fig. 1b). Furthermore, functional annotations allow us to separate sparse driver mutations from confounding passenger mutations. The end result is the aggregation of mutations that are individually low in frequency into high-frequency regulatory recurrence events, based on their convergence into common target genes (Fig. 1c). This can be achieved even in the absence of clusters of contiguous alterations, protein coding or otherwise.

**Figure 1.**
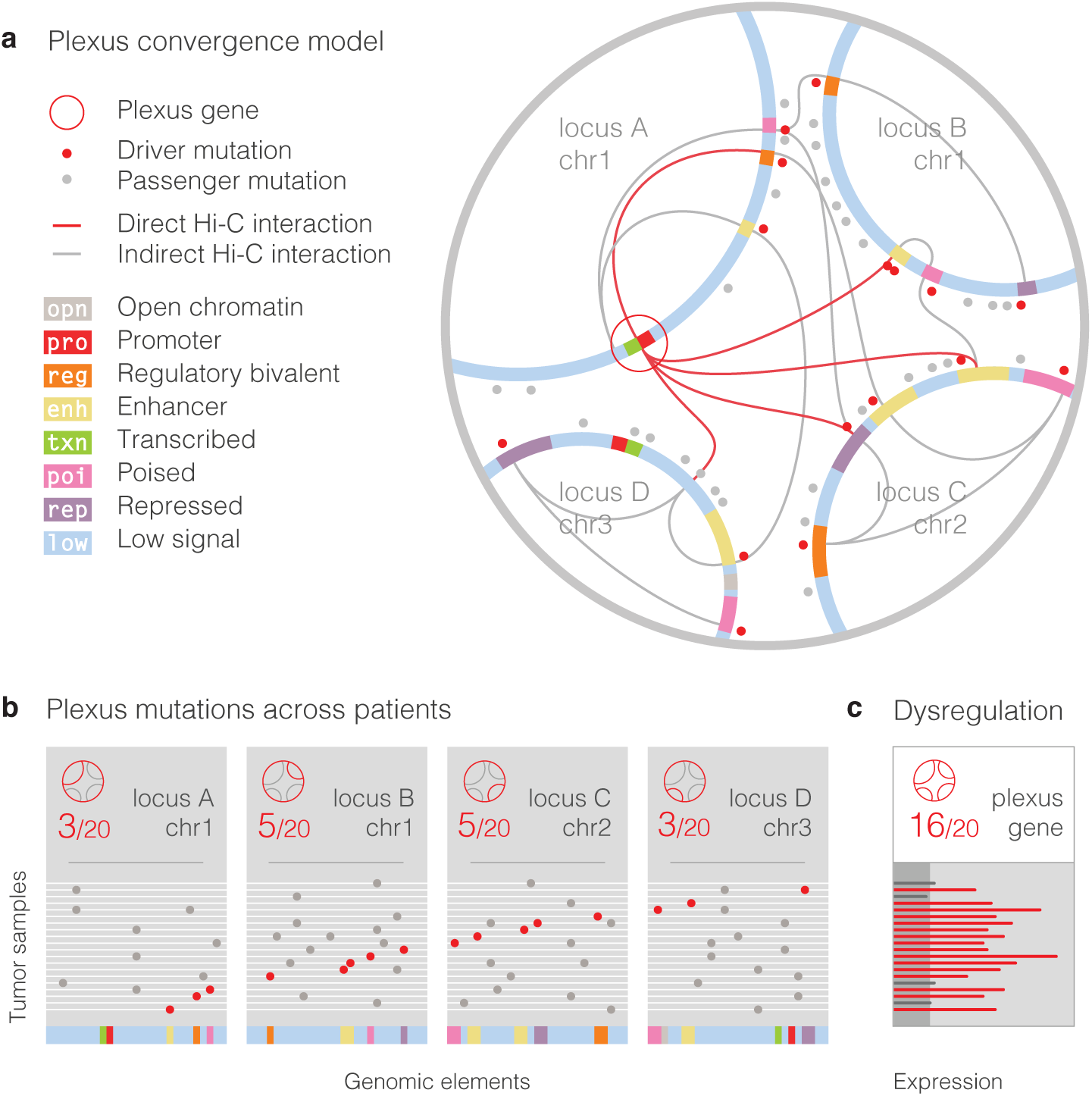
The plexus framework reveals the convergence of dispersed and heterogeneous mutations **a**, Visualization of a hypothetical plexus composed of four loci. The four loci contain a variety of active and inactive genomic elements (arcs with colored segments within an encompassing gray circle). The plexus’ gene (red circle) has an active promoter (‘pro’, red) and a transcribed region (‘txn’, green). Intra- and inter-chromosomal Hi-C interactions connect the plexus’ gene to regulatory elements elsewhere in the plexus (red Bezier curves). Additional interactions help maintain the four loci together within the volume of the cell nucleus (gray Bezier curves). Dispersed mutations in active elements (putative drivers, red dots) are interspersed with mutations in inactive chromatin (likely passengers, gray dots). Active elements harboring mutations include enhancers (‘enh’, yellow) and poised regulatory elements (‘poi’, pink), among others. **b**, Dispersed active (red) and interspersed inactive (gray) mutations arranged by tumor sample. Mutational heterogeneity across samples leads to low-frequency mutation events at single elements or loci, each accounting for a common mechanism of tumorigenesis in only a small number of patients (3-5 out of 20). **c**, Aggregating low-frequency mutation events through a gene’s plexus can reveal high-frequency driver events through convergent dysregulation of the same gene; a single mechanism now accounting for a majority of patient tumor samples (16 out of 20).

## RESULTS

### Plexus assembly from matched ChIP-seq and Hi-C

We first establish the prostate-specific plexus of every protein-coding gene in the human genome. Regulatory annotations are highly tissue specific (Lupien et al. 2008). We therefore use the RWPE1 prostate cell line as a reference, and profile five histone modification marks using ChIP-Seq. We use ChromHMM (Ernst and Kellis 2012) to define eight chromatin state annotations consisting of: promoters (‘pro’) with strong H3K4me3 but no H3K4me1; enhancers (‘enh’) with H3K4me1 but no H3K4me3; regulatory elements (‘reg’) marked with both enhancer and promoter signatures; transcription-associated regions (‘txn’) with H3K36me3; poised elements (‘poi’) with H3K27me3 and at least one other active mark; repressed elements (‘rep’) with H3K27me3 only; and low-activity regions (‘low’) where no marks are detected (Fig. 2a, S1). Raw plexi exhibit an over-abundance of active regulatory states (‘opn’, ‘pro’, ‘reg’, ‘enh’), especially through proximal interactions (Table S1). We treat open chromatin regions (‘opn’), based on DNaseI in RWPE1 (ENCODE Consortium 2012), as a separate class, regardless of their enclosing chromatin state.

**Figure 2.**
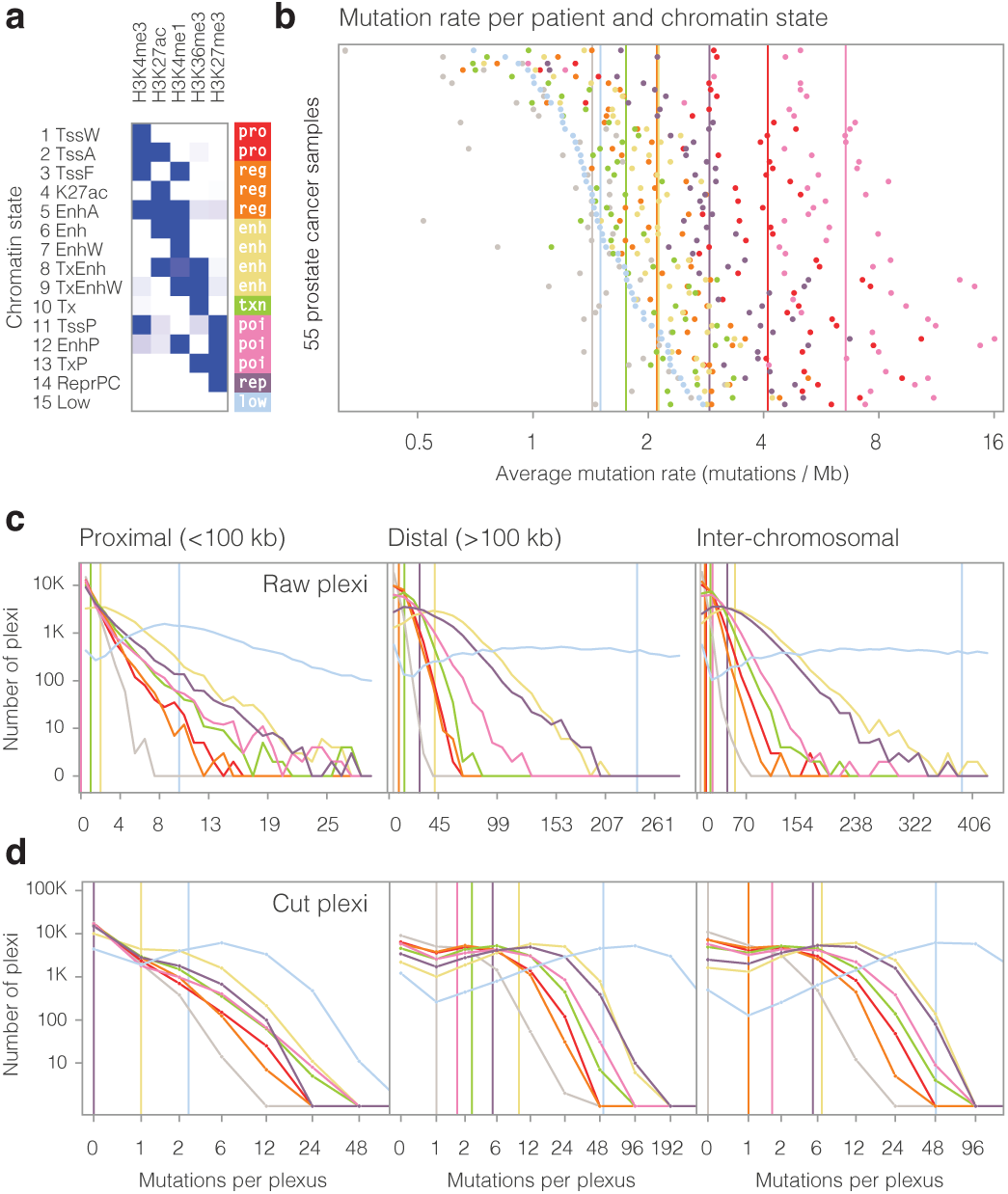
Plexus assembly by connecting dispersed regulatory mutations to protein-coding genes using matched ChIP-seq and Hi-C **a**, Chromatin states in normal prostate. We profiled five histone marks (columns) in RWPE1 cells, and used ChromHMM to learn 15 chromatin states (rows), which we further group in eight aggregate states (colors). We treat open chromatin regions, regardless of the enclosing chromatin state, as a separate class (not shown). **b**, Mutation rate heterogeneity across chromatin states and tumor samples. Scatter plot showing mutation rates (x-axis) across 55 prostate tumor samples (y-axis) for eight chromatin states (colors). Tumor samples are sorted by average mutation rate in low-activity regions. Colored vertical bars indicate median mutation rate for each state across tumor samples. **c**, **d**, Linear histograms show the number of plexi (y-axis) by the mutation count in connected elements (x-axis) for raw (c) and cut plexi (d) assembled around all protein-coding genes. Plexus mutation counts are calculated separately for chromatin states (colors) and three classes of distance to the plexus gene (columns). Colored vertical bars indicate the median number of mutations per gene for each chromatin state.

We find large variation in mutation rates, across both chromatin states and tumor samples (Fig. 2b). Open chromatin regions show the lowest mutation rate (1.5 mutations/Mb), consistent with previous reports (Stamatoyannopoulos et al. 2009; Polak, Querfurth, and Arndt 2010; Polak et al. 2014), attributed to their increased association with the DNA repair machinery (Liu, De, and Michor 2013). However, even outside DNaseI regions, mutation rates vary greatly across chromatin states (from 1.6 to 6.9 mutations/Mb on average), and across tumors (from 0.7 to 2.8 mutations/Mb for low-activity regions). Mutation rates do not correlate with GC content, CpG dinucleotide rate, nucleotide, or di-nucleotide composition (Table S2), and chromatin states preserve their relative mutation rates across tumors (Fig. 2b), suggesting sequence-independent mechanisms, possibly due to differential interactions with the repair machinery by chromatin regulators. Surprisingly, once open chromatin regions are excluded from chromatin state annotations, the expected inverse correlation between epigenomic activity and mutation rate is lost. Instead, promoter regions free of open chromatin show the second highest mutation rate.

We link these regulatory annotations to each gene using a prostate-specific map of chromosome interactions that is also derived from the RWPE1 cell line (Rickman et al. 2012) **(Data S1)**. With this data we generate two types of plexus for each protein-coding gene: a raw plexus, which contains any locus with evidence of interaction, and a cut plexus, in which each interaction is assessed using a permutation test and filtered based on a p-value cutoff of 0.05 **(See Methods)**. While the short lengths of regulatory elements make contiguous, single-element recurrence rare (Fig. S2), through its plexus a gene can be associated with a much higher richness of variation across a patient cohort. The set of raw plexi associates genes with a median of 106 linked proximal elements (within 100 kb), 1420 distal intra-chromosomally (*cis*), and 1824 inter-chromosomally (*trans*) interacting elements; providing an abundant source of mutation to each gene (Fig. S3a, data S2). Indeed, each gene interacts with a median of 21 mutations in proximal elements, 399 mutations in distal *cis*-elements, and 625 mutations in *trans*-interactions (Fig. 2c), providing sufficient power to study plexus-level mutation. We use the raw plexi to identify recurrent driver events across the 55 patients; in this way we cast a broad net that we progressively tighten to further concentrate our candidates. Conversely, cut plexi associate each gene with a median of 30 interacting proximal elements (within 100 kb), 319 distal cis-elements, and 227 trans-elements (Fig. S3b, data S2). This leads to a more focused source of mutation, with each gene being associated with a median of 1 mutation in proximal elements, 24 mutations in distal *cis*-elements, and 17 mutations in *trans*-elements (Fig. 2d). We exploit the stringent interactions of the cut plexi for intra-chromosomal interactions to look for enrichment of distal mutations in cancer-dysregulated genes.

### Dysregulated genes enrich in plexus mutations

To study the relevance of plexus mutations as a mechanism of gene dysregulation we use whole transcriptomes obtained from 16 of the 55 patients with whole genomes (Baca et al. 2013). Every tumor sample has a matched normal sample from adjacent prostate tissue. From the normal tissue, we establish a range of expression for every gene and use each distribution to normalize tumor transcriptomes (Fig. 3a, see Methods). Tumor samples showed significantly greater variance (Wilcoxon P < 10^-15.7^) in expression than normal prostate samples (Fig. 3b). For each gene, we search for pairs of tumor samples where one gene instance is dysregulated in one patient and the other instance is unchanged. Pairing multiple gene instances in this manner allows us to compare a large set of dysregulated gene instances against a control set that is matched one-to-one so as to preserve the genomic and epigenomic properties between the two sets (Fig. 3a). Mutational properties between the two sets, which contain different compositions of patients, are corrected by incorporating the patient- and chromatin-specific mutation rates into the enrichment calculations (See Methods). Additionally, we only use dysregulated and unchanged gene instances when the normal prostate samples for the same two individuals show normal expression. We do this to ensure that dysregulation is not already present in the adjacent matched tissue before tumor development. Through this approach we identify 17,850 dysregulated-unchanged paired samples over a total of 2,579 genes (Fig. 3c, data S3), of which 83% are up-regulated (14,893), and only 17% are down-regulated (2,957).

**Figure 3.**
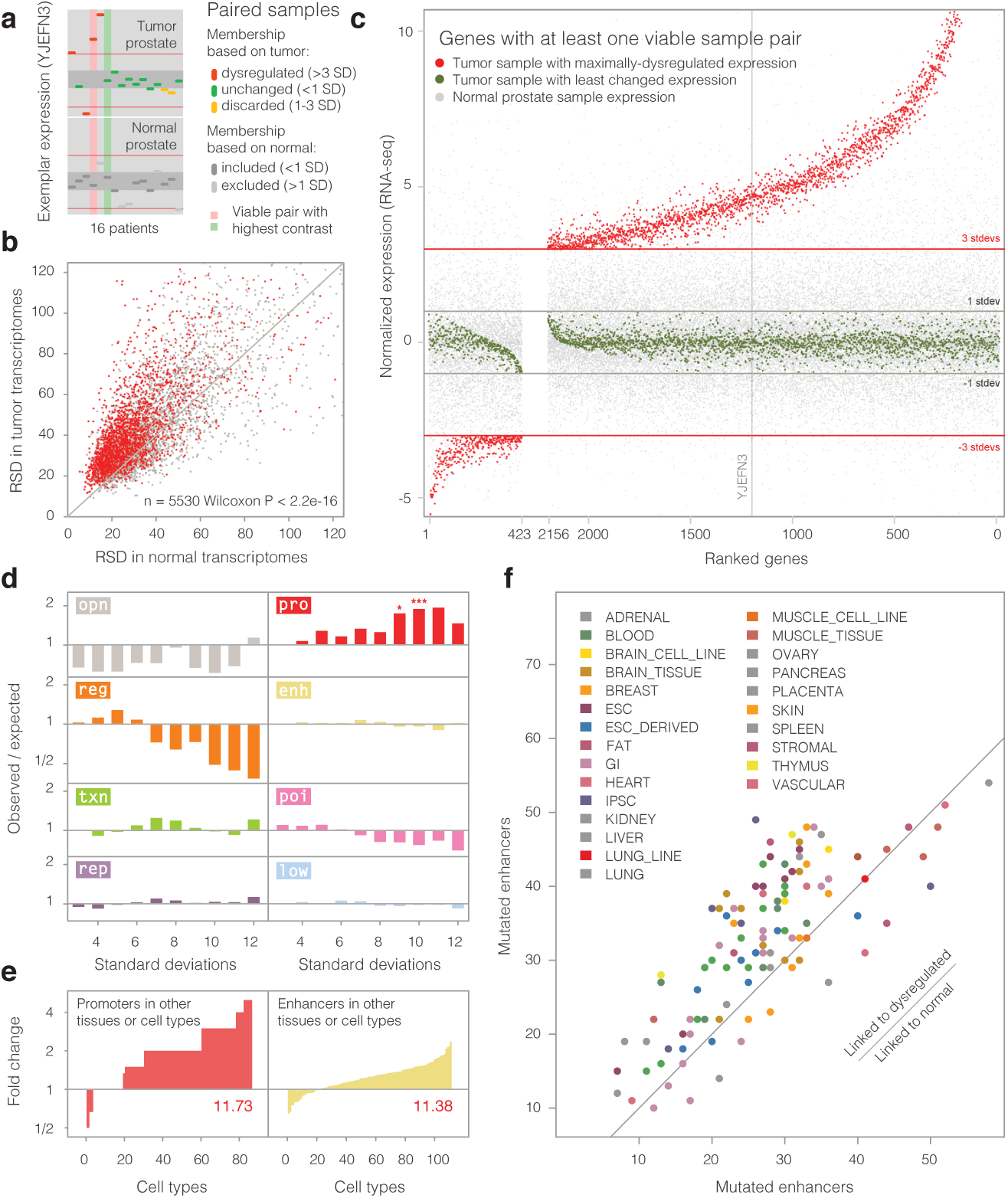
Dysregulated genes are enriched in dispersed plexus mutations with regulatory activity or latent regulatory potential **a**, An example of a dysregulated and unchanged gene instance pair (normalized expression; y-axis) across 16 patients (x-axis) for exemplar gene YJEFN3. Tumor (top) and matched normal samples (bottom) are used to identify viable pairs across the whole transcriptome. A viable pair (red and green columns) is composed of a dysregulated (top, red, >3 SDs) and an unchanged tumor sample (top, green, <1 SD) where both samples have normal expression (bottom, dark grey, <1 SD) in the matched normal prostate tissue. Gray regions and red horizontal lines indicate the +/-1 and +/-3 SD intervals. **b**, Gene expression in tumors is more variable than in normal tissue. Relative standard deviation (RSD) in 16 prostate tumor samples (y-axis) and 16 matched normal prostate samples (x-axis). Genes selected as dysregulated in viable gene instance pairs (panels a, c) are shown in red. **c**, Dysregulated and unchanged gene instance pairs are predominantly up-regulated. Scatter plot shows the normalized expression values (y-axis) for every gene used in the analysis (x-axis). Viable pairs and controls are plotted for every gene, as defined in panel a. **d**, Enrichment scores for mutations linked to dysregulation. Histograms show the log2 ratio of observed over expected mutation counts (y-axis) at increasing levels of dysregulation (x-axis; P_Bonf_<0.05; 20,000 permutations). **e**, Mutations in low activity regions linked to dysregulated gene instances enrich in promoters (red) and enhancers (yellow) of non-prostate tissues. Histograms showing the ratios of overlap between mutations linked to dysregulation against those linked to unchanged gene instances (fold changes; y-axis) across Epigenome Roadmap cell and tissue types (x-axis; Wilcoxon P = 10^-11.73^ and 10^-11.38^) **f**, Scatterplot showing the number of Epigenome Roadmap enhancers overlapped by mutations linked to dysregulation (y-axis) against those linked to unchanged gene instances (x-axis) for mutations in low-activity regions in prostate (Wilcoxon P = 10^-4.81^).

We test the hypothesis that the plexi of dysregulated gene instances are enriched for mutations that are distal to the gene (>100 kb from gene body). For this we only use high confidence interactions from the cut plexi (Permutation P < 0.05; intra-chromosomal, See Methods), and restrict our analysis to up-regulated genes where we have more power to detect an effect. We perform enrichment tests for mutations across all chromatin states and over a range of magnitudes for up-regulation. We find a consistent enrichment for ‘pro’ elements that increases as dysregulation becomes more extreme (Fig 3d). This enrichment peaks at 10 standard deviations from normal expression (Permutation P_Bonf_ < 0.05). These enrichments remain even when we repeat the analysis removing all gene instances with copy number alterations, albeit with less statistical power (Fig. S4). Interestingly, ‘opn’ and ‘reg’ elements show signs of being protected from mutation; perhaps they represent a set of highly optimized activators that are less likely to increase their function through random mutation.

Based on our previous work regarding the gain and loss of enhancers in tumor initiation in colon cancer (Akhtar-Zaidi et al. 2012) and tumor progression in breast cancer (Magnani et al. 2013), we hypothesize that a fraction of mutations in ‘low’ elements for prostate might be active in non-prostate cell lines, thus driving dysregulation activity in prostate cancer through out-of-context de-repression of existing but dormant regulatory elements, as opposed to creating them from scratch. Indeed, ‘low’ elements harboring mutations are strongly enriched for both promoter (Wilcoxon P < 10^-12^) and enhancer states (Wilcoxon P < 10^-11^) in other cell types (Fig. 3e). These are active in a diverse panel of cell and tissue types (Roadmap Epigenomics Consortium 2014), including immune cells, GI-tract, and ESCs (Fig. 3f), suggesting co-option of diverse, non-prostate elements.

### Plexus recurrence test reveals hidden drivers

Having established that dysregulated genes are enriched in distal plexus mutations, we next sought to identify individual genes with an excess of mutations in their plexi in the whole genomes of the 55 tumor samples. It is highly unlikely that positive selection acts exclusively on cancer driver genes through coding and proximal promoter mutations, especially considering how most GWAS variants that increase the risk of cancer act through distal regulatory mechanisms, and the bewildering diversity of mutational processes in tumor evolution.

A plexus recurrence test is faced with the same confounders as recurrence tests for coding genes, namely mutational heterogeneity across genomic regions due to cell-type-specific transcriptional activity (Lawrence et al. 2013) and chromatin states, in addition to variability among patient and tumor mutational signatures. In testing a plexus we must also account for changes in the confounders across the constituent loci (Fig. 4a). Incidentally, we find that regional mutational heterogeneity is even more extreme than previously recognized, following a power-law distribution at the 50 kb scale (Fig. 4b), suggesting localized mutational bursts. The plexus recurrence tests starts by gathering a plexus’ mutations, regional mutation rate estimates and chromatin states for all the loci it contains. These layers of information are stored at a resolution of 100 bp; we refer to these intervals as ‘tiles’. We retrieve a tile array for every protein-coding gene in the human genome through that gene’s raw plexus. The tile array is then decomposed into the two major confounders: regional mutation rate and chromatin state (Fig. 4c). We compress mutation rates into 15 bins of exponentially increasing mutational intensity (taken from the tile’s 50 kb context). Chromatin states are stored as the 8 categories previous described.

**Figure 4.**
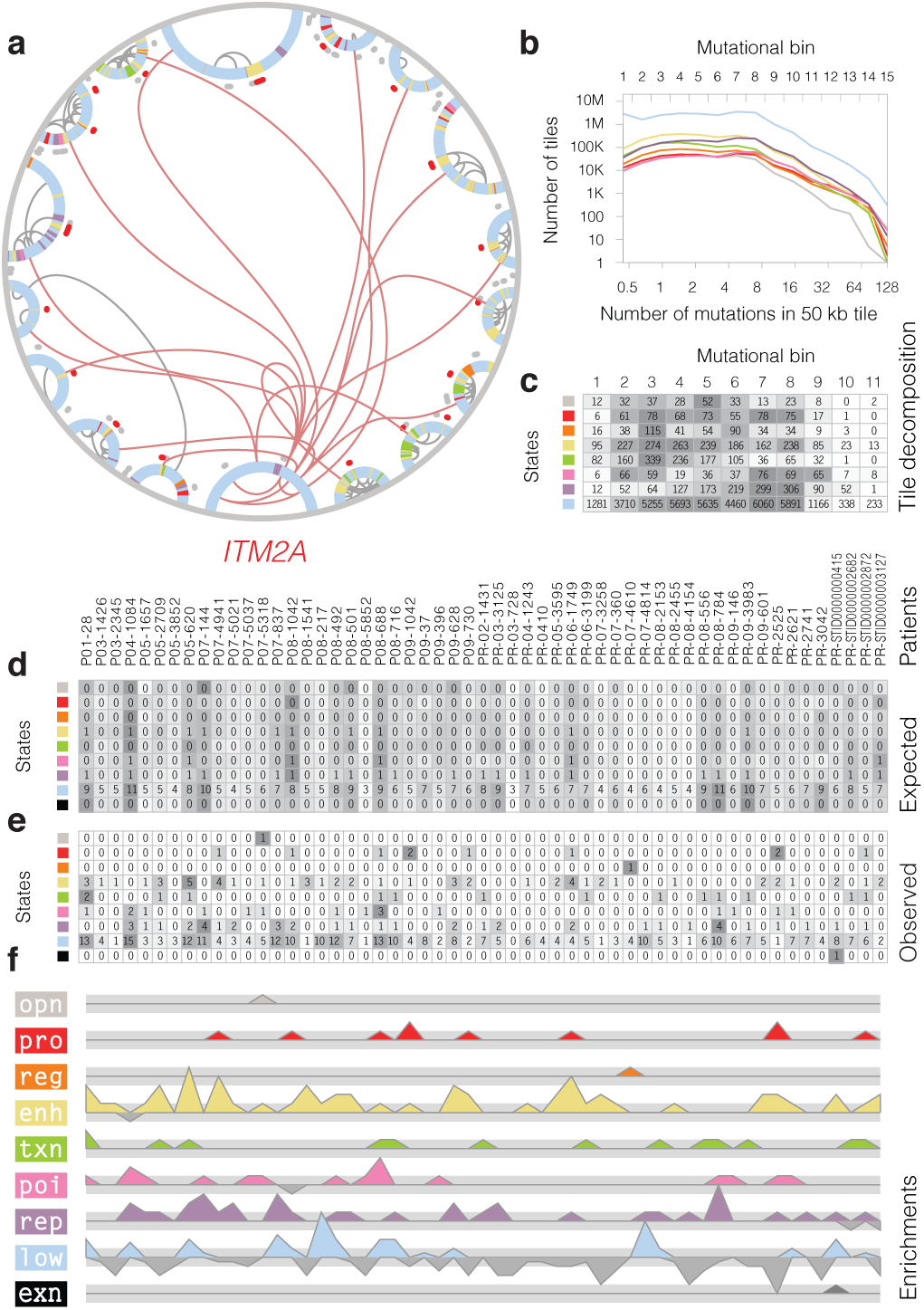
Plexus recurrence test identifies putative driver genes while controlling for mutational heterogeneity across chromosome regions, chromatin states, and patients The recurrence test for the *ITM2A* plexus. **a**, The *ITM2A* plexus, visualized as in Fig. 1a, is used as an example. **b**, Regional mutation rate heterogeneity. Mutation rates (x-axis) assessed over a sliding window of 50 kb for every 100 bp tile in the human genome (number of tiles, y-axis) follow a power-law distribution. The top axis shows the boundaries of the mutational bins used in tile resampling. **c**, First, we tally the number of 100 bp tiles in the *ITM2A* plexus and store them in a matrix structured by mutational bin (columns) and chromatin state (rows). This is the plexus tile decomposition for *ITM2A*. **d**, Second, we randomly sample 100 bp tiles from the whole genome in such a way as to match the tile decomposition of the *ITM2A* plexus. From this we obtain a matrix of expected mutation counts for every patient (columns) and chromatin state (rows) combination. **e**, Third, we tally the mutations in the *ITM2A* plexus and obtain the matrix of observed mutation counts. **f**, Fourth, we use the observed and expected counts to obtain enrichment scores for each patient and chromatin state combination. Finally, the enrichment scores are aggregated across patients in order to obtain a final, permutation-based p-value for each chromatin state. Shading denotes intensity in the matrices; rows are independently normalized.

Statistical significance is computed through permutation. The tile decomposition of a plexus is used to guide the random sampling of tiles from the whole genome so as to match the chromatin state and regional mutation rate properties of the test plexus. We then retrieve patient mutations for the permuted tile array so as to match the heterogeneity of the mutation rate in the patients. By aggregating mutations over all permutations we obtain the expected number of mutations for each patient and chromatin state over the tile array (Fig. 4d). The expected mutation counts allow us to convert observed mutation counts (Fig. 4e) into enrichment scores (Fig. 4f). Because the enrichments we previously observed for dysregulated genes in plexus mutations are highly dependent on chromatin state, we test each chromatin state separately. We combine the enrichment scores across all 55 patients to obtain a final list of p-values for each of the chromatin states and for the gene’s exons (Table S3). We refer to this procedure simply as ‘the plexus recurrence test’ (See Methods).

Applying the plexus recurrence test to the 55 prostate cancer whole-genome sequences, we identify 15 recurrently mutated plexi that are statistically significant (Table S3, data S4). The genes varied greatly in enriched chromatin state (‘txn’, ‘pro’, ‘rep’, ‘poi’, ‘enh’), patients harboring mutations (35%-89%), number of mutated elements (7-62), and number of mutations (24-150). The plexi do not share regulatory regions, however, the *RRAD* plexus also contains the *FAM96B* gene (Table S4). These 15 genes lie in a small number of common pathways, suggesting higher-order functional convergence. They are involved in cell growth, migration and proliferation through overlapping roles in androgen, insulin and circadian rhythm signaling (*INSRR*, *PLCB4*, *CRY2*, *RRAD*, *SPANX* and *SSX*), immune evasion (*ITM2A*, *IDO2*, *ZC3H12B* and *ZBED2*), mitochondrial function (*COQ3* and *SLC25A5*) and vascularization (*EDNRA*). Two genes remain uncharacterized (*C14orf180* and *ZCCHC16*). Several of these genes have been linked to cancer: *INSRR* (Hua et al. 2013), *RRAD* (Reynet and Kahn 1993), *SSX* (D’Arcy et al. 2014); and prostate cancer, specifically: *CRY2* (Chu et al. 2007; Zhu et al. 2009). However, the most clinically relevant gene may be *IDO2*, a partner of IDO1 (Metz et al. 2014). IDO genes constitute a key mechanisms of immune evasion and have recently become central targets in immuno-oncology (Sheridan 2015).

### *PLCB4* plexus 3C loops and epigenomic landscape

Having identified a list of 15 candidate driver plexi, we select one for experimental validation. To do this, we use the p-values assigned to each locus of the raw plexi that were obtained through permutation. We then apply increasingly stringent thresholds to each plexus and recompute the plexus recurrence test at every step. As the cut plexi shrink they lose both mutated and non-mutated loci, making the recurrence signal oscillate and ultimately decay completely. The *PLCB4* plexus has the most robust recurrence signal among all 15 plexi (Fig. 5a). The signal comes from 79 mutations in ‘txn’ elements that are distal to the *PLCB4* gene body. Of these, 30 are on the same chromosome and spread over 12 loci. We perform chromatin conformation capture (3C) experiments and confirm 5 out of the 12 interactions with the *PLCB4* promoter (Fig. 5b, S5, Table S5, See Methods). We group these elements into four loci and refer to them by their mega base coordinates on chromosome 20: 9.0 (which contains the *PLCB4* gene body), 9.2, 10.4, 18.5 and 30.0. In addition to the 3C experiments, the five loci are woven together by numerous direct and indirect Hi-C interactions (Fig. 5c).

**Figure 5.**
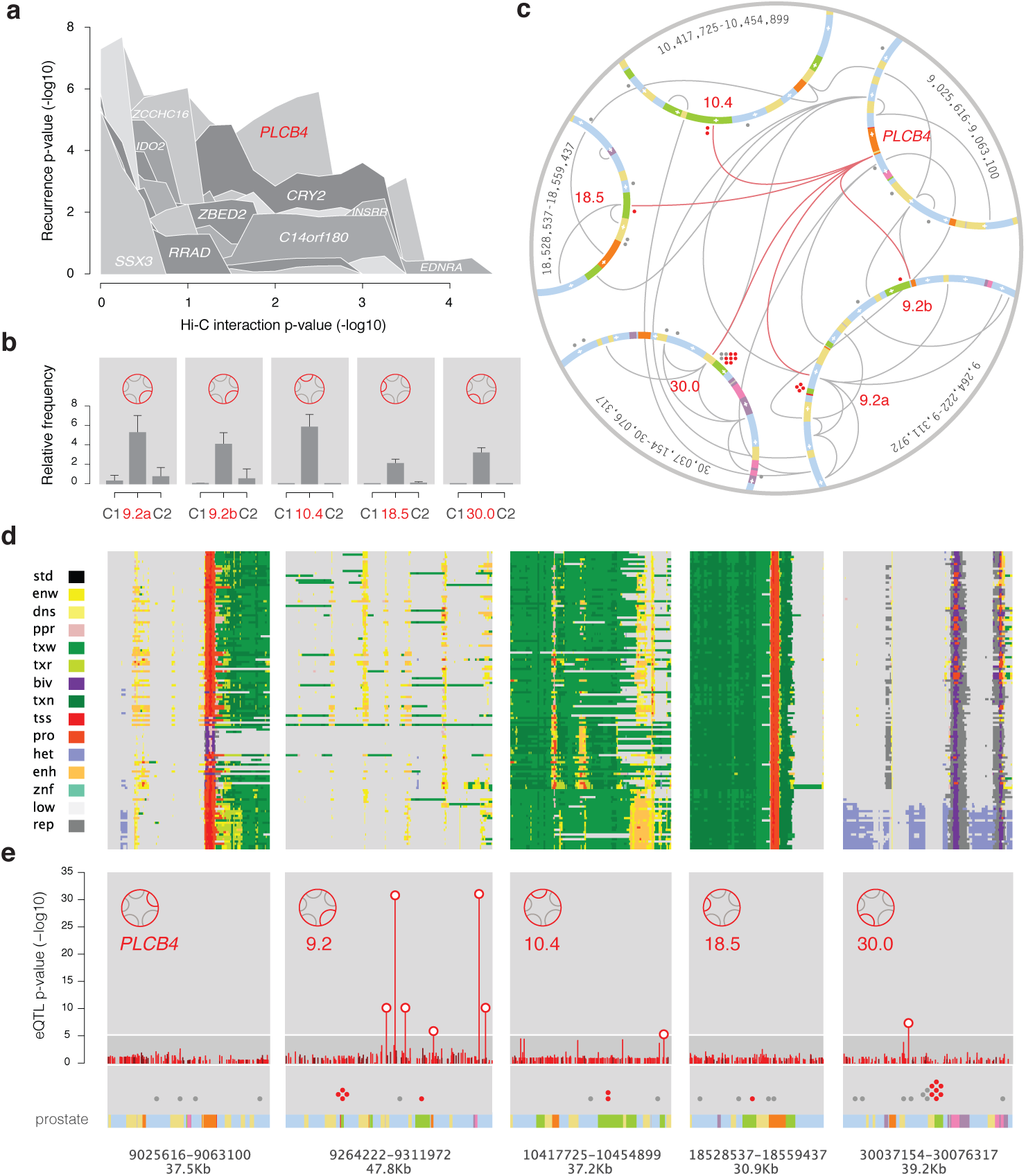
*PLCB4* plexus signal robustness, 3C structure, epigenomic landscape and eQTLs **a,** Signal decay rate of the plexus recurrence test for the top 15 plexi. We re-compute p-values (y-axis) as we increase the stringency of Hi-C interactions (y-axis). The *PLCB4* plexus (red) shows the most robust signal. **b,** Chromatin interactions between the *PLCB4* promoter and the rest of the plexus in the RWPE1 prostate epithelial cell line. Bar plots show the chromatin conformation capture (3C) interaction strength between the *PLCB4* promoter and each of the other four loci in the plexus. Adjacent *Eco*RI fragments are used as controls. Error bars represent SD of three biological replicates assayed in duplicate. **c,** The 3C-validated *PLCB4* plexus visualized as in Fig. 1a with the following differences: only mutations for the significant ‘txn’ state are highlighted in red, scale marks for each locus are drawn every 5 kb, and Bezier curves depict 3C-validated (red) and Hi-C (grey) interactions. **d,** Epigenome Roadmap annotations for the five loci of the 3C-validated *PLCB4* plexus (x-axis) depicted linearly across 127 cell types (y-axis). **e,** GTEx eQTLs for *PLCB4* computed from 87 prostate samples. Scatter plot depicts the –log_10_ p-value (y-axis) of *PLCB4* eQTLs for 783 variants (97, genotyped, red; 686 imputed, dark red) contained in the five loci of the 3C-validated *PLCB4* plexus (x-axis).

The 3C-validated loci contain 15 mutations for 12 patients in the significant chromatin state, ‘txn’, and 36 mutations for 24 patients over all chromatin states (Fig. 5c). The significant (‘txn’) and extended (all states) mutation sets represent 22% and 44% of the 55 total patients, respectively. The recurrence frequency at the *PLCB4* plexus is high compared to coding genes identified in exome sequencing studies of prostate cancer. The top three genes by frequency, *SPOP*, *TP53* and *PTEN*, are mutated in 13%, 6% and 4% of patients, respectively (Barbieri et al. 2012). However, the impact of mutations in coding regions is well understood. Annotations across 127 reference epigenomes from the Roadmap and ENCODE projects (Ernst and Kellis 2015; Roadmap Epigenomics Consortium 2014; ENCODE Consortium 2012) help infer the regulatory potential of the loci mutated in the *PLCB4* plexus. Gene dysregulation through out-of-context de-repression would require latent or poised regulatory elements to be present in the loci in cell types other than prostate. Although incomplete, the Roadmap and ENCODE collection of cell types can give some indication of regulatory activity, even when highly specific to a few non-prostate cell types. The annotations show highly diverse regulatory contexts in the ∼40 kb regions containing the mutations (Fig. 5d). Some loci show consistent activity across all tissues, whereas others reveal striking prostate specificity. Finally, three out of the four distal loci in the *PLCB4* plexus (9.2, 10.4 and 30.0) contain eQTLs for *PLCB4* expression based on the analysis of 87 prostate samples from the Genotype-Tissue Expression (GTEx) project (GTEx Consortium 2015) (Fig. 5e, data S5, See Methods).

### *PLCB4* plexus disruption and the PI3K pathway

The study of cancer recurrence in coding regions benefits from our knowledge of the genetic code and allows for the filtering of mutations based on synonymity. In the regulatory setting we need tissue-specific annotations and models of protein-DNA binding to obtain a similar understanding. Unfortunately, a comprehensive regulatory code is still unavailable. However, we are able to infer a portion of the code in prostate cells by leveraging a subset of transcription factors binding profiles across canonical prostate cell lines. We first scanned the 15 mutations in the *PLCB4* plexus for binding of transcription factors in any human cell type using ReMap (Griffon et al. 2015). Five of the seven mutations at the 30.0 locus overlap binding sites for ERG, TP63, SP1 or BRD4 (Fig. 6a). All of these factors are involved in gains, losses or fusions in prostate cancer (Sankpal et al. 2011; Tucci et al. 2012; Tomlins et al. 2005; Asangani et al. 2014). We then interrogated the five core histone marks (H3K4me1, H3K4me3, H3K36me3, H3K27me3 and H3K27ac) and open chromatin in five additional prostate cell lines, ranging from normal tissue (RWPE1, PWR1E) to low (LNCaP), moderate (DU145) and high (22Rv1, PC3) tumorigenicity (Fig. 6b). The reference for this study, RWPE1, shows active transcription marks across all four loci in addition to promoter marks at 9.2 and 30.0. The 9.2 and 10.4 loci show variability in the normal tissue and loss of histone marks across all cancer cell lines. Locus 18.5 is consistently active in all cell lines, whereas locus 30.0 shows a strong gain in enhancer marks in cell lines with high tumorigenicity and one normal cell line. These results suggest that gains and losses of activity in the *PLCB4* plexus relate to a range of cancerous phenotypes in a standard collection of prostate cell lines.

**Figure 6.**
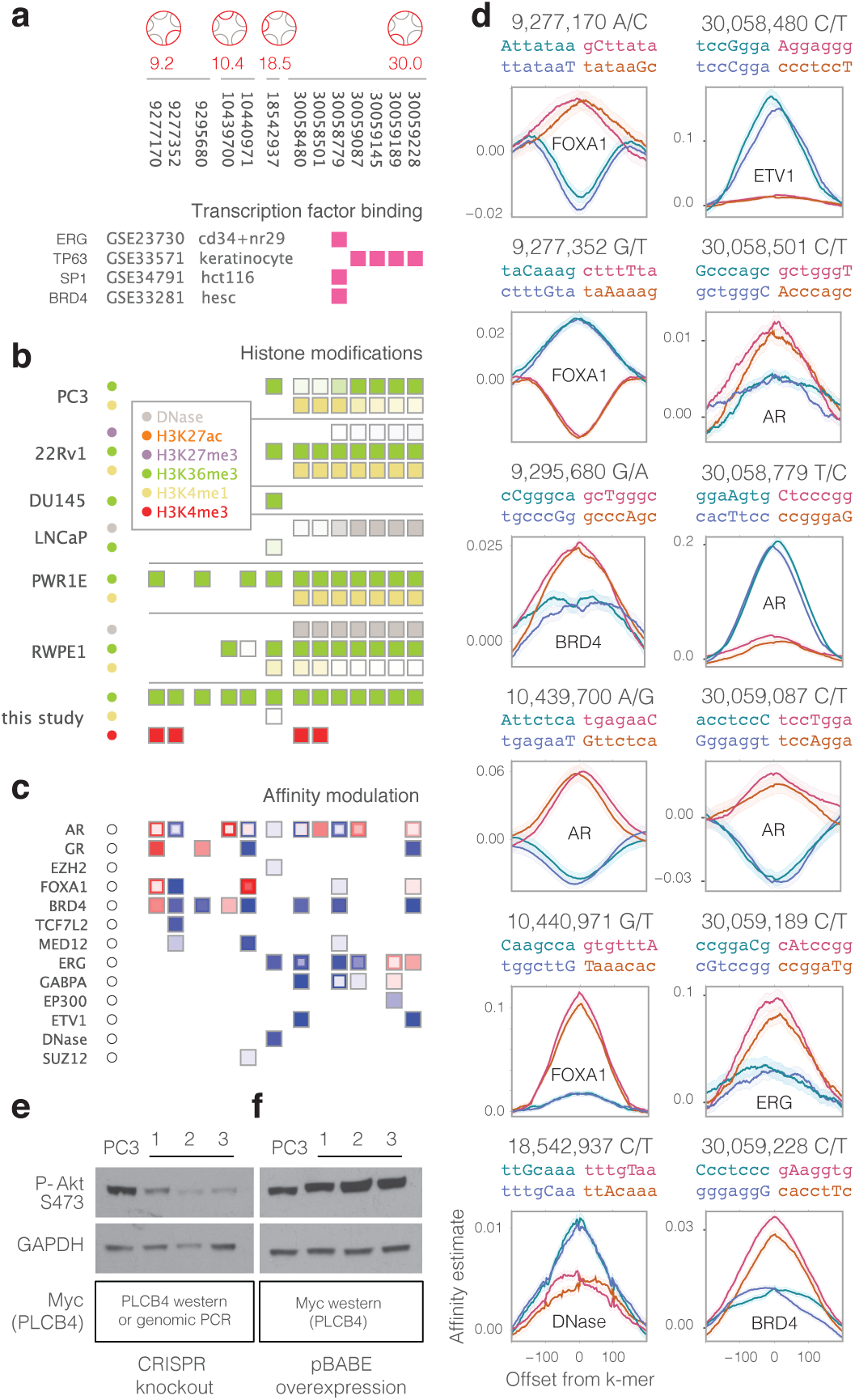
*PLCB4* plexus mutations affect the binding of key prostate transcription factors and PLCB4 disruption affects the PI3K pathway Regulatory annotations for 15 non-coding mutations (columns) in the 3C-validated *PLBC4* plexus overlapping ‘txn’ state elements. **a**, Cancer transcription factor binding sites in non-prostate cells overlap mutations in the 30.0 locus. **b**, Histone marks in six prostate cell lines of increasing tumorigenic potential (from the bottom to the top) show losses at the 9.2 and 10.4 loci, but consistent activity at the 18.5 and 30.0 loci. Distance of the mutation to the closest region is indicated by saturation, where full saturation indicates direct overlap and minimum saturation indicates a 10 kb distance. **c**, Intra-genomic Replicates (IGR) measures of affinity modulation for a collection of prostate-related transcription factors assayed in prostate cell lines. Gains (red) and losses (blue) in binding are shown for significant results. Nested cells show results from multiple replicates of the same factor. **d**, IGR profile plots showing the most disruptive effect for each of the 14 single-nucleotide mutations in the 3C-validated *PLCB4* plexus. Each plot shows the affinity estimation (y-axis) for each of the normal (blue) and mutated (red) sequences. Affinity estimate profiles are shown over a 400bp window (x-axis) centered on the k-mer. Only the maximum affinity k-mer for each allele is presented in the final IGR result. **e**, *PLCB4* loss lowers PI3K signaling. Three independent *PLCB4*-deficient PC3 lines were engineered using CRISPR/Cas9 and their lysates immunoblotted with the indicated antibodies. **f**, *PLCB4* overexpression increases PI3K signaling. Three independent PC3 lines stably transduced with pBABE myc-PLCB4 were isolated and their lysates immunoblotted with the indicated antibodies.

In order to assess the role of the tumor mutations in the *PLCB4* plexus on these activity patterns, we consider their effects on the binding of key transcription factors that are likely mediating the deposition of active histone marks. We built affinity models for all prostate transcription factors in the Cistrome database of ChIP-seq experiments (http://cistrome.org/db) using the Intra-Genomic Replicates (IGR) method (Cowper-Sal·lari et al. 2012) **(See Methods)**. We found that mutations in the 9.2 locus tend to increase the binding of AR, GR, FOXA1 and BRD4 factors, whereas mutations in the 30.0 locus tend to consistently decrease binding of BRD4, ERG, GABPA and ETV1 (Fig. 6c). However, the most commonly affected factor is AR, with disruptive mutations in all four loci. Each of the 14 mutations probed with IGR (single-nucleotide) had a large effect for at least one of the factors. Among the most dramatic results we find the creation of a FOXA1 binding site at locus 10.4 and the destruction of two binding sites for AR and ETV1 at the 30.0 locus (Fig. 6d). Interestingly, we found a significant enrichment in binding-altering mutations for ETV1 (QFDR < 0.024) across all 35 mutations at the *PLCB4* locus **(See Methods).**

PLCB4, or phospholipase Cβ4, has been extensively studied in the context of circadian rhythms and auriculocondylar syndrome, where it has strong effects when disrupted (Park et al. 2003; Rieder et al. 2012). The role of PLCB4 in prostate cancer is unknown, although it has been identified in a set of 96 genes associated with in vivo progression to castration-recurrent prostate cancer (Romanuik et al. 2010). Considering its ability to directly affect cell membrane lipid metabolism and the phosphatidylinositides, we tested the impact of *PLCB4* deletion or overexpression on the phosphoinositide-regulated PI3K/AKT signaling pathway in PC3 cells. While *PLCB4* deletion led to a decrease in PI3K/AKT signaling throughput (Fig. 6e), its overexpression resulted in activation of the PI3K pathway (Fig. 6f), revealing a direct link between the levels of *PLCB4* expression and the activity of this major oncogenic signaling cascade.

## DISCUSSION

We present the first scan for driver genes with a plexus recurrence test and, more generally, demonstrate the use of long-range chromatin loops to decipher genetic heterogeneity by collapsing the combinatorics of high-order, multi-locus interactions with the three-dimensional structure of the genome. We find that dispersed non-coding mutations that are individually too low in frequency for viable statistical analysis nevertheless converge into high frequency recurrence events. These events reveal novel driver genes with known and putative roles in prostate cancer even in the absence of proximal mutations (protein coding or otherwise), which have been the focus of previous studies. Furthermore, these genes show pathway-level convergence in androgen, insulin and circadian rhythm signaling, immune evasion, mitochondrial function and vascularization, providing new insights into the biological processes known to underlie prostate cancer. Most notably, we identify *PLCB4*, which we validate as capable of affecting the canonical PI3K cancer pathway, and *IDO2*, a gene whose function is currently a central target in immuno-oncology and responsible for hundreds of millions of dollars in biotech investment and acquisitions (Sheridan 2015).

We believe the plexus framework will be especially valuable for cancers with low mutation rates. Ependymoma, an extreme example, lacks any detectable mutational recurrence (Mack et al. 2014). We can now qualify this statement by saying that it lacks proximal and contiguous mutational recurrence for any gene. However, plexus mutational recurrence might still be an important driver in such tumors. Indeed, many of the genes we have identified have no mutations in their gene bodies or in proximal regions. The plexus framework might also shed light into the temporal and functional interplay of diverse types of mutations through tumor initiation and progression. We hypothesize that the first somatic aberrations that propel a cell towards cancer act much like risk-associated variants. These primordial DNA mutations or epigenomic alterations are likely distal and regulatory in nature, having subtle and heterogeneous effects at first. But as these changes accumulate, triggering out-of-context de-repression of regulatory elements, they gradually converge on the hallmark pathways of cancer. Furthermore, plexus mutations are less likely to be immunogenic, allowing for the creation of a neutral evolutionary space in which pre-cancerous cells would accumulate a high degree of variability which, in turn, would provide a rich substrate for selection when it arises. Exploring this ‘plexus first’ hypothesis of cancer emergence and evolution poses a daunting challenge, as it will require the comprehensive mapping of multi-locus interactions in normal and cancerous cells across numerous human tissues.

Beyond cancer, the plexus framework introduced here is broadly applicable to the analysis of common, rare and private variants in any human disease or trait. Genetic heterogeneity in sequence association studies currently hinders our ability to uncover the molecular basis of heritability in complex traits. We hope that applying the plexus framework to association studies will reveal trait-associated plexi (and the genes therein) where, although each constituent locus explains only a small subset of affected individuals, the whole plexus accounts for a significant proportion of the cohort. The low frequency problem of contiguous variation has led researchers of type 2 diabetes to conclude that rare variants do not contribute significantly to disease risk (Fuchsberger et al. 2016). However, with our method the frequency of collapsed alleles increases and the number of tests decreases. Therefore, the plexus framework might boost power in such studies and allow for a more effective use of whole genome sequence data. Similarly, the plexus framework could constitute a broader foundation for the interpretation of individual genomes. Instead of limiting analyses to the 1.5% of the genome that encodes proteins, all variants would be used. Ultimately, we wish to help advance personalized therapeutics and precision medicine by empowering those using whole-genome sequencing in the understanding, prediction and treatment of complex disease.

## AUTHOR CONTRIBUTIONS

RCS led the study and conceived of the plexus framework in discussions with JHM. RCS, JHM, NASA, ML and MK developed the project. RCS and NASA designed, implemented and ran all computational analyses. SLE and JDF validated all interactions for the *PLCB4* plexus. VS, ML, JH and KJK elucidated the role of PLCB4 in the PI3K pathway. RCS, NASA, and MK made the figures. RCS and MK wrote the manuscript with the help of all authors.

## ACKNOWLEDGEMENTS

We thank Levi Garraway, Sylvan Baca, Franco Preparata, Cigall Kadoch, Aviad Tsherniak, Melina Claussnitzer, Angela Yen, Robert Altshuler, Pouya Kheradpour, Avanti Shrikumar, Nezar Abdennur and Wouter Meuleman. This work was supported by NIH grants LM009012 and LM010098 to JHM and R01 HG004037 to MK, NHMRC project grant 1058415 to SLE and JDF, Prostate Cancer Canada Movember Rising Star award RS2014-04 to ML, the NSF CAREER award 0644282 to MK, and the Stanford Graduate Fellowship to NASA.

## METHODS

### Data sources

Normal prostate Hi-C chromosome interaction data (chromatin loops) in the RWPE1 cell line were downloaded from the Gene Expression Omnibus (GEO) database at www.ncbi.nlm.nih.gov/geo (accession number: GSE37752)(Rickman et al. 2012). Prostate cancer-normal whole genome and transcriptome pairs for 55 prostate adenocarcinoma patients were obtained through the database of Genotypes and Phenotypes (dbGaP) at www.ncbi.nlm.nih.gov/gap (accession number phs000447.v1.p1) and directly at the Broad Institute (Baca et al. 2013). Gene annotations were downloaded from GENCODE at http://www.gencodegenes.org (version 18). DNase annotations generated as part of the ENCODE project were downloaded from the UCSC Genome Browser at genome.ucsc.edu. Epigenome Roadmap promoter and enhancer annotations were obtained at the Broad Institute (www.broadinstitute.org/∼meuleman/reg2map/HoneyBadger_release).

Normal prostate transcriptomes and genotypes for eQTL analysis were obtained from the Genotype-Tissue Expression (GTEx) project at www.gtexportal.org. Additional ChIP-seq data for epigenomic and transcription factor binding analyses were downloaded from cistrome.org/db. Copy number data for the 55 prostate adenocarcinoma patients was obtained from www.cbioportal.org.

### Epigenomic profiling of normal prostate chromatin state

Healthy prostate ChIP-seq data for five core histone marks (H3K4me1, H3K4me3, H3K36me3, H3K27me3 and H3K27ac) and DNase was generated in the RWPE1 cell line with the same protocols used in Cowper-Sal·lari et al. (Cowper-Sal·lari et al. 2012) except using sonication instead MNase and the Illumina HiSeq 2000 instead of the Genome Analyzer. Chromatin states were learned with ChromHMM (Ernst and Kellis 2012). We used 100 bp elements in order to maintain the granularity of the smaller DNase annotations from ENCODE. The Epigenome Roadmap 15-state ChromHMM model was further aggregated into eight broader functional categories: open chromatin, promoter, regulatory (with mixed promoter and enhancer marks), enhancer, transcribed (with no other function), poised (promoter, enhancer and transcribed), repressed, and low-activity regions (no marks). These aggregate states are denoted by the following three-character mnemonics: opn, pro, reg, enh, txn, poi, rep and low (Fig. 1, S1). All source data generated by the Lupien lab can be downloaded from www.pmgenomics.ca/lupienlab/tools.html.

### Plexus assembly

#### Objective

The plexus framework seeks to identify cellular functions that are dysregulated through alterations distributed over multiple loci in the context of cancer recurrence and trait association studies. It is predicated on the notion that the set of loci that affect any given cellular function are likely to be non-contiguous and sparse on the one-dimensional sequence of the genome. Identifying these sets of loci from sequence alone or through exhaustive testing is currently infeasible; we therefore use experimentally derived annotations for both the locations and interactions of these sets of loci in order to address the sparseness and non-contiguity problems, respectively. Furthermore, both the locations and interactions of the loci that determine cellular functions are highly variable from one cell type to another. The use of cell types that are relevant to the trait or disease under study is critical. In principle, the framework can use any source of alteration. In this study we focus on somatic, single nucleotide variants in a cohort of 55 prostate adenocarcinoma patients, but we look forward to expanding the repertoire of alterations to germline variants, all structural classes of mutations, and epigenetic and transcription factor binding changes. We define the plexus as the comprehensive set of genomic loci that when altered can modulate a cellular function. In this study, we focus on the expression of protein-coding genes. We first build the cell-type-specific plexus for every protein-coding gene; this includes regulatory elements that are both proximal and distal to the gene body, on both the same (*cis*) and other (*trans*) chromosomes. Specifically, we do this with histone marks and chromosome interactions (chromatin loops) obtained for the same cell-type in which the mutations originate. Ultimately, this allows us to search for genes that have more mutations in their regulatory elements or gene body than expected by chance alone. We use two types of plexus in this study: a lenient raw plexus, which intends to encompass as many of the true interacting loci as possible, and a stringent cut plexus, where interactions are filtered based on a permutations test.

#### Raw plexus

We retrieve transcription start site (TSS) and exon annotations from the GENCODE database (version 18) for all proteincoding genes. We segment the human genome (hg19) into 100 bp tiles and assign a chromatin state to every tile. Chromosome interactions were originally generated through the Hi-C sequencing technique. Interactions between loci are mediated through DNA binding proteins. Each binding site at the terminus of an interaction likely covers a few tens of bases. However, the Hi-C technique can only resolve the positions of the interaction termini to a few kb. This is due to the use of the *Hin*dIII restriction enzyme that cuts the DNA. The ends of Hi-C sequencing reads are unambiguously assigned to *Hin*dIII fragments, but the exact location of the terminus within the fragment cannot currently be determined. We refer to each segment of the genome contained between two *Hin*dIII restriction sites as an anchor. Every 100 bp tile is therefore contained within a *Hin*dIII anchor, and anchors are connected to each other through Hi-C interactions. Through this network we assign tiles to the plexus of every gene. First, we retrieve all anchors within 10 kb up and downstream of the gene’s TSS. For each anchor at the TSS, we then retrieve all other anchors connected through Hi-C interactions. Each of these anchors is extended 10 kb in both directions. We then store all tiles that overlap either a TSS anchor, any of the gene’s exons, or any of the extended anchors at the distal ends of the Hi-C interactions. The list of 100 bp tiles is then sorted and filtered down to an array of unique tile indices. Raw plexus tile arrays frequently span multiple chromosomes and several Mb (Fig. S3). They also contain a variety of chromatin states and disparate regions of the genome with highly discordant mutation rates. The plexus tile array is the representation of all the proximal and distal elements that potentially impinge on a gene’s function. We compute plexus tile arrays for every protein-coding gene in the human genome **(Data S1),** and then retrieve tumor mutations for every array **(Data S2).** The raw plexi constitute the basis for our plexus recurrence test.

#### Cut plexus

Several factors can account for the presence of reads spanning two loci in a Hi-C library, many of which are confounders to identifying true regulatory interactions. We set out to assign a measure of confidence on putative interactions present in each of the raw plexi previously defined. The cut plexus program takes the raw plexus of a gene as input. It then loads the Hi-C matrices for the chromosome that contains the gene. The matrices are binned at 10 kb intervals for the intra-chromosomal interactions (inter-chromosomal are discarded for the cut plexi) and computed using the Mirny lab’s hiclib library (mirnylab.bitbucket.org/hiclib). The raw data from Rickman et al. contains four replicates: GFP1, GFP2, ERG1 and ERG2 of varying sequencing depth. Interactions are tested separately across the four replicates and aggregated at the end of the program’s execution. The GFP and ERG conditions are intended to simulate normal and cancerous cell states. We want to capture interactions that can occur across the carcinogenic continuum. Furthermore, we are primarily interested in identifying loci that can interact rather than loci that interact frequently. Therefore, because these loops might vary over time, identifying a loop in a single replicate is still of interest. We then assess each of the locus pairs in the raw plexus. Interactions that are less than 100 kb from the gene promoter are tagged as “proximal”. Because for now we are only concerned with intra-chromosomal interactions, the rest are tagged as “distal_cis”. We generate a p-value for each of the four replicates by permuting matrix counts between the two loci.

#### Hi-C null

-This null distribution represents the expected counts given the distance between loci and the genomic background at each of the loci. To compute the null we start by tallying the observed matrix count. This number is the largest value among the cells corresponding to the bins containing the pair of loci or any of the eight adjacent cells. We do this in order to include arrangements in which the probed loci and interaction anchors are separated in adjacent bins. The anchors that mediate the interactions between genes and regulatory elements might not overlap perfectly (Bailey et al. 2015). True interactions can be mediated by physical interactions that are some kb away. Additionally, restriction fragment processing makes the exact location of these interactions uncertain. We then take a segment of the diagonal that crosses the cell addressed by bin coordinates of the loci. This diagonal slice of the matrix expands 100 cells upstream and downstream parallel to the diagonal and two cells perpendicular. We tally the matrix values (balanced reads) over the diagonal slice and then randomly reassign the same number of reads over a matrix of the same dimensions as the diagonal slice. We count the number of times that we see values larger than or equal to the observed matrix count in the permuted matrix. We then regenerate the randomized matrix 100 times. The p-value for each replicate is calculated by dividing the number of cells with tallies larger or equal than the observed value over the total number of cells and matrix randomizations. We combine the p-values over the four replicates using Fisher’s method. This final p-value is what we use to filter the locus pairs in the raw plexus to generate cut plexi of varying stringency. Through this approach we attempt to estimate the probability of the observed read count given the genomic context of the two loci. Our intention is to correct for the interface or area of interaction between the loci at the level of chromosome territories. We do this at a scale that is much larger than the individual loci but that retains the local conformational background. We run the permutation test on all GENCODE V18 genes marked as ‘protein_coding’ for all chromosomes except Y and M, which yields a total of 20,233 genes. We use a p-value threshold of 0.05 to define all subsequent statistics for cut plexi.

### Dysregulated gene analysis

#### Core scores

-We identify dysregulated instances of genes in specific patients across 16 cancer transcriptomes by normalizing every gene’s expression using an aggregate of values in the matched normal prostate tissue. The normalized measure we use is a slight variation on z-scores, which we refer to as “core scores”. We calculate the median and standard deviation (SD) of each gene from the 16 normal transcriptomes. Because prostate cancer has a strong genetic component we consider that some gene instances might already be dysregulated in the matched normal tissue. Therefore, we discard the six most extreme values for each gene. We do this by sorting all gene-instances for each gene and finding the sequence of ten consecutive instances with the smallest SD. Because removing the extremes of any distribution will warp the estimates of its SD, we estimate the distortion factor extracting the core from random sets of 16 values sampled from the normal distribution and correct the core-scores accordingly (distortion factor ∼2). Finally, we filter out lowly expressed genes where the normal core has a median FPKM value under 0.3 (Ramsköld et al. 2009).

#### Matching of pairs

-We search for genes where at least one instance has an absolute core score larger than three SDs (dysregulated) and one other instance has an absolute core score smaller than one SD (unchanged). These dysregulated and unchanged gene instance pairs allow us to study transcriptional dysregulation in one patient while having a control instance of the same gene in a second patient. The expression values for patients in normal tissue need to have an absolute core score smaller than one SD in order to be included in the analysis. We do this to ensure that dysregulation is not already present in the adjacent matched tissue before tumor development. We identify 17,850 viable, dysregulated and unchanged gene instance pairs over a total of 2,579 genes (Fig. 3c, data S3), of which 83% are up regulated (14,893), and only 17% are down regulated (2,957).

#### Enrichment scores

-We use the cut plexi (p-value < 0.05) to assign mutations to each of the gene instances for all viable pairs. We use cut plexi and intra-chromosomal interactions in order to enrich for true interactions, as the Hi-C is notoriously noisy. Because we are strictly interested in the effect of distal mutations, we only consider those that are beyond 100 kb of the plexus’ gene promoter. Additionally, we restrict our analysis to up-regulated genes where we have more power to detect an effect. Having gathered mutations for both dysregulated and unchanged gene instances in the tumor samples, we can compute the enrichment of mutations in dysregulated gene instances by comparing the two groups in aggregate. Because the specific contribution of different classes of regulatory elements to tumorigenesis is unknown, we compute enrichment scores for each of the chromatin states separately. Pair resampling -We test several intensities of dysregulation for enrichment in mutations, from core scores between 3 and 12 SDs. For each baseline, we pick all the pairs in the 17,850 viable pair set where the core score of the dysregulated gene instance exceeds or equals the baseline core score. A permuted set of pairs of the same size as the one defined by the core score baseline is resampled within the same collection of genes by randomizing the indices of the dysregulated and unchanged instances between genes. This is similar to randomly sampling 16x16 patient sample pairs, but with a non-uniform distribution over the patients. By simply permuting the indices between genes, the distribution approximately preserves the patient proportions among the dysregulated and unchanged categories across all genes. We then retrieve the plexus mutations for the dysregulated and unchanged gene instances and append them to their respective mutation matrices (gene instances by chromatin states). As we increase the core score baseline, the number of viable pairs decreases and the analysis loses power.

#### Patient balancing

-Patient proportions in the dysregulated and unchanged matrices are not balanced. When selecting gene instance pairs for a given core score baseline, we tally the number of times each patient appears in either category. Based on the mutation rate heterogeneity across patients, we compute the expected dysregulated to unchanged ratio of mutations for each chromatin state. We then compare this to the dysregulated to unchanged ratios between the two mutation matrices. The ratio of ratios is log transformed and constitutes the enrichment score for each chromatin state at each baseline. We derive a p-value for this test by counting the number of times that the observed enrichment score is larger than the enrichment scores in the permuted gene instance pairs. Finally, we correct the p-values by the 10 core scores baselines and 8 chromatin states tested (Bonferroni; 80 tests).

#### Copy number correction

-If dysregulation is due to regional copy number alteration, then the region is likely to have more mutations assigned to it during variant calling, as the copies are conflated in the variant calling. This could lead to the appearance of enrichment when compared to an unchanged gene instances in a patient with no copy number alterations in that region. We therefore add an additional condition to the previous approach in order to avoid the possible effect of copy number alterations in our study of dysregulated genes. When collecting gene instance pairs for each core score baseline, a pair is only considered if it has copy number data available and neither gene instance has a copy number alteration. This filter is applied to both the observed and permuted pair sets.

### Plexus recurrence test

#### Tile resampling

-The plexus recurrence test is designed to identify gene plexi that harbor more mutations than are expected by chance alone. It estimates the expected number of mutations through resampling. Its null distribution accounts for critical confounders that have been previously identified in the search for driver events in cancer genomes (Lawrence et al. 2013). Namely, it accounts for heterogeneous mutation rates across patients, chromatin states and genomic regions. The null distribution is computed from a resampling matrix in which all 100 bp tiles used to assemble the plexi are binned based on their chromatin state and regional mutation rate. The regional mutation rate at each tile is calculated from its surrounding 50 kb by pooling mutations across all patients. We find that larger windows fail to account for the broad variations in mutation rates, whereas smaller windows are similar in size to genes, which can be legitimate units of selection. Each tile is assigned its mutational bin number based on the log of its regional mutation rate (Fig. 4b). The two-dimensional binning of all tiles in the human genome results in a matrix of tile arrays of 15 mutation bins by 8 chromatin states. Each tile retains the assignment of mutations for each patient. Thus, preserving the mutation rate heterogeneity present in the cohort. A similar two-dimensional binning is applied to the tile array of the test plexus, where instead of tile index lists we store tile tallies (Fig. 4c). We refer to this matrix of tile tallies as the tile decomposition of the plexus. It encodes our understanding of the mutational confounders across the loci in the plexus; the attributes that affect the accumulation of mutations but that are independent of selection. The plexus tile decomposition guides the resampling in each permutation. Tiles are picked at random and with replacement from the resampling matrix so as to match the tile decomposition of the test plexus and control for the mutational confounders. The collection of permuted tile arrays constitutes the null distribution with which to estimate expected mutation rates for each patient and chromatin state combination.

#### Centroid score

-In the next step, the plexus recurrence test retrieves the patient mutations for the test plexus and each of the permutation tile arrays. A tile is considered mutated for a patient if it contains one or more mutations. Each tile is therefore associated with a binary array with an entry for each patient. Mutations for the permuted tile arrays retain their original assignments to patients in order to control for mutation rate heterogeneity in the cohort, including patient-specific variation in chromatin sate mutation rates. Observed mutations in the test plexus tile array are compared to mutations in the permutation tile arrays using a separate centroid score for each chromatin state and the exon annotations contained in the plexus (referred to just as states for brevity, but now including a ninth category for exon elements). The centroid score is computed in the following manner in order to allow for aggregation across samples and states. Mutations are tallied in state by patient matrices for all tile arrays (Fig. 4d, e). Tally matrices for the permutations are stored as a three-dimensional null volume. Each list of mutation tallies for each state and patient combination is upper quartile normalized across the permutation dimension. Positive normalized tallies summed across patients constitute the centroid score, which are calculated separately for each state. This produces nine centroid scores for the test plexus and each of the permutations. Each patient contributes to the centroid score proportionally to how much it positively deviates from what is expected for that patient and chromatin state combination (Fig. 4f).

#### Significance

-Statistical significance for the test plexus is computed by comparing its centroid scores against those of the permutations. The size of the null distribution (number of permutations) is increased until a reliable p-value is determined. Only plexi in which one or more of the nine tests (eight chromatin states and exons) have two or fewer permutations that have a centroid score larger or equal to that of the null pass on to the next round. We perform nine tests on each of the 20,318 protein-coding genes and correct the p-values accordingly with the Benjamini-Hochberg method **(Data S4).**

#### Discussion

-Many features remain to be incorporated and explored in future version of the plexus recurrence test. First, due to limited availability and high cost of Hi-C data, we construct the plexus of each gene in reference to a single prostate cell line. However, profiling chromatin states, DNase hypersensitivity regions, and especially Hi-C interactions in each tumor and normal sample individually would provide for a much richer analysis of each patient’s disease. The direct incorporation of variability in plexus structure between individuals and the regulatory rewiring within tumors would be particularly interesting. Second, even though we corrected for mutation rate differences between chromatin states, we treated all mutations in the same chromatin state as equally likely to have a regulatory effect and drive tumorigenesis. Protein-coding models that distinguish between synonymous and non-synonymous mutations and attempt to predict the effect of variants are readily available. However, similar models for regulatory alterations are not as developed. Richer regulatory models will allow future iterations of the test to incorporate the magnitude and direction of effect for non-coding mutations in the expectations derived from resampling. For example, by pre-computing the likelihood of a mutation perturbing enhancer activity or modulating binding of a transcription factor. The models will eventually leverage tumor-specific information on regulator activity, such as the intra-cellular concentration of aberrant transcription factors. Third, the three-dimensional structure of the genome has been shown to be scale-free, with rich patterns of chromosome interactions at multiple scales. Therefore, our test of non-contiguous recurrence should not be limited to a single scale, in this case the organization of regulatory elements around a single gene, but should consider both smaller and larger structures. The tested units could range from enhancer clusters or sets of interaction termini to transcriptional factories or topological domains. The pathway-level convergence we observe among the 15 drivers we identify suggests another set of layered recurrence tests across the hierarchy of cellular functions. A hierarchical recurrence test could reveal convergent mutations in the merged plexi of protein complexes, metabolic pathways and cancer hallmarks. Finally, the plexus recurrence test should combine somatic and germ line variants of strong and weak effects, that are rare to common in frequency, both protein-coding and non-coding, to cover the entire causal timeline of cancer, from inherent risk to emergence and evolution. This would constitute the first approximation of a unified model of tumorigenesis

### Chromatin conformation capture (3C) of the *PLCB4* plexus

The RWPE1 (normal prostate epithelial) cell line was kindly provided by Dr Jyotsna Batra (Queensland University of Technology, Brisbane, Australia) and cultured in KSFM supplemented with 5ng/ml epidermal growth factor, 25μg/ml bovine pituitary extract and 2mM glutamine. 3C libraries were generated using *EcoRI* as described previously (Ghoussaini et al. 2014). 3C interactions were quantitated by real-time PCR (Q-PCR) using primers designed within *EcoRI* restriction fragments (Fig. S5, Table S5). Q-PCR was performed on a RotorGene 6000 using MyTaq HS DNA polymerase (Bioline) with the addition of 5 mM of Syto9, annealing temperature of 66°C and extension of 30sec. 3C analyses were performed in three independent library preparations with each experiment quantified in duplicate. Bacterial artificial chromosome (BAC) clones were used to create artificial libraries of ligation products in order to normalize for PCR efficiency. Q-PCR products were electrophoresed on 2% agarose gels, gel purified and sequenced to verify the 3C product.

### eQTL analysis of variants in the *PLCB4* plexus

The Genotype-Tissue Expression (GTEx) project is a publicly available resource that provides matched genotype and expression data from normal human donors (GTEx Consortium 2015). 87 of these donors have transcriptome data (RNA-seq) for prostate, in addition to genotype and covariates data. We intersect the five 3C-validated loci with the GTEx genotypes and obtain 783 variants (97 genotyped, 686 imputed) for the *PLCB4* plexus. Nine transcripts of *PLCB4* (ENSG00000101333.12) have detectable expression in at least one of the 87 prostate transcriptomes: ENST00000278655.4; a, ENST00000334005.3; b, ENST00000378473.3; c, ENST00000378501.2; d, ENST00000416836.1; e, ENST00000464199.1; f, ENST00000473151.1; g, ENST00000482123.1; h, ENST00000492632.1; i.

We compute expression quantitative trait loci (eQTLs) for the nine transcripts and the 783 variants using the Matrix eQTL R package (Shabalin 2012). The Bonferroni cutoff for statistical significance is 10^-5.2^ (0.05 / (783 * 9)). We used the age, race and ethnicity of the GTEx donors as covariates in the analysis.

### Intra-genomic replicates (IGR) analysis of *PLCB4* plexus

We downloaded signal tracks from Cistrome DB for 118 ChIP-Seq experiments performed in prostate cell lines, as well as ERG and GABPA in Jurkat cells (Sharma et al. 2014). We construct 400 bp window 7mer and 8mer IGR models for each track, as previously described (Cowper-Sal·lari et al. 2012) except that we do not apply the competition filter. We use 373,359 active prostate elements genome wide as the IGR regional filter. We define these regions as the union of peaks across 29 experiments profiling DNase, H3K27ac, H3K4me1, H3K4me2, and H3K4me3 in the same prostate cell lines. Using these models, we run the IGR algorithm on 14 out of the 15 mutations in ‘txn’ states and 35 of the 36 total mutations contained in the *PLCB4* plexus (only single-nucleotide variants). Many of these mutations show statistically significant affinity modulation of transcription factor binding between the reference (germline) and alternate (tumor) alleles (Bonferroni corrected). IGR filters -We have updated the original IGR program with two new filters. First, in order to discard noisy affinity models lacking sufficient instances of a given k-mer genome wide to make a clean prediction we devise the “quality” and “symmetry” filters. IGR computes an averaged binding profile for every k-mer; in this case along a 400 bp window centred on the k-mer. Every k-mer has two profiles for its forward and reverse complement orientations, and every IGR result has two final k-mers for the highest affinity among the reference and alternate allele k-mer sets. The correlation between the forward profile and the mirror image of the reverse profile constitutes the measure of quality. The correlation between the forward profile and its own mirror image constitutes the measure of symmetry. We then remove any mutation results where either reference and alternate final k-mer profiles had either symmetry or quality smaller than 0.5 and both had symmetry and quality smaller than 0.85. Second, in order to only select mutations for which the effect size was large enough we calculated baseline-offset affinities for each of the final k-mer profiles. This measure compares the affinity centred at the k-mer, minus the average of the signal 195-200 bp away from the k-mer in both forward and reverse orientations. Using these, we define the “maximum prominence” as the highest absolute baseline-corrected affinity in either the reference or alternate allele within 200 bp of the k-mer and the “maximum difference” as the largest absolute difference between baseline-corrected reference and alternate alleles within 200 bp of the k-mer. We exclude all mutation results for which the ratio between the maximum difference and maximum prominence was less than 0.5.

#### Enrichment analysis

-We test *PLCB4* plexus mutations for enrichment of affinity modulating results that satisfy all of the previous filters. We assess the set of mutations in the *PLCB4* plexus as a whole using Fisher’s exact test within each experimental setup. Enrichments were corrected using FDR and only significant IGR mutations were used when counting (q-value < 0.024; estimate = 6.73 for 8-mers)

#### *PLCB4* overexpression

PC3 cells were transfected with PolyJet (SignaGen Laboratories) as per manufacturer’s instructions. Briefly, cells were plated the night before transfection to achieve 70% confluence on 10 cm dishes. PC3 cells were transfected with empty vector, pcDNA3.1-mycHis-PLCB4, p3xFlag-CMV10-PTEN or both. Forty-eight hours later, the cells were harvested by washing and scraping in ice-cold PBS followed by centrifugation at 1,500*g* for 5 minutes at 4°C. The cell pellet was lysed in 160 μL of CHAPS lysis buffer (40 mM HEPES pH 7.5, 0.3% CHAPS, 120 mM NaCl, 1 mM EDTA, 50 mM sodium fluoride, 20 mM beta-glycerophosphate, protease inhibitor cocktail) on ice for 20 minutes. The cell lysate was clarified by centrifugation at 15,000*g* for 15 minutes at 4°C and the supernatant was normalized for total protein using the Bradford assay (Biorad). SDS loading buffer was added to the normalized samples and 30 μg of total protein was loaded on 8% acrylamide gels and transferred to PVDF membranes. The membranes were blocked in 5% BSA in TBST and immunoblotted with anti-PLCB4 (sc-20760, Santa Cruz Biotech), anti-phospho Akt (#4058, Cell Signaling), anti-phospho Erk (#9106, Cell Signaling), anti-Akt (#4691, Cell Signaling), anti-Erk (#9102, Cell Signaling) and anti-PTEN (#9559, Cell Signaling).

## SUPPLEMENTARY FIGURES

**Figure S1.**
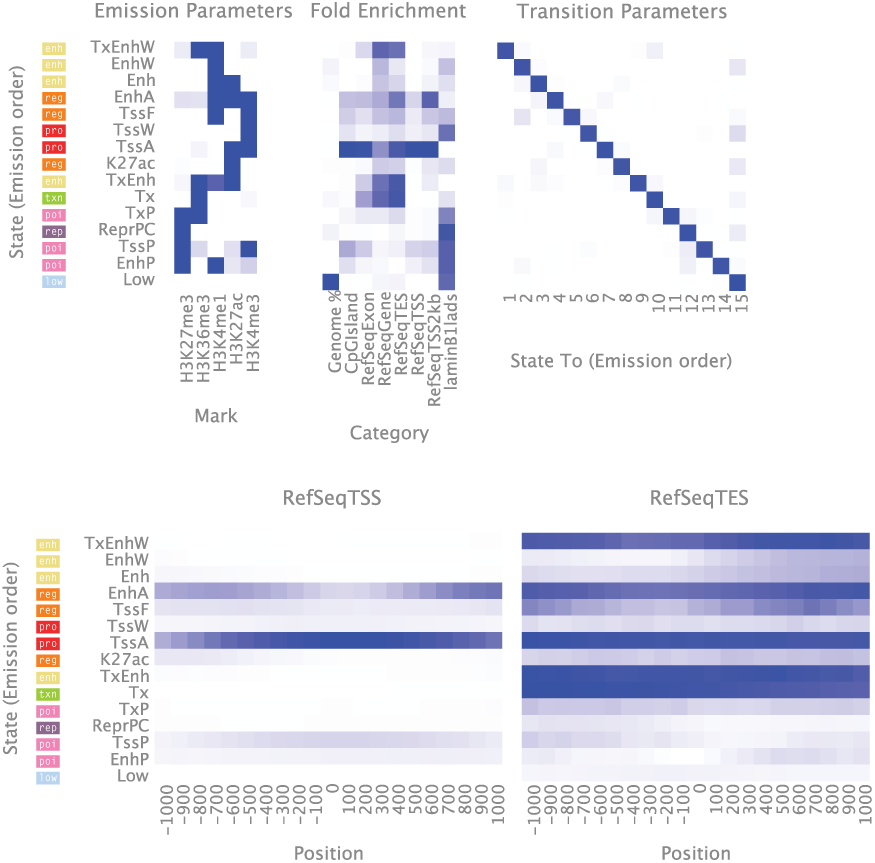
ChromHMM states and aggregate states. Emission parameters for the 15-state model learnt on the five core histone marks. Aggregate states represent simplified regulatory roles.

**Figure 2.**
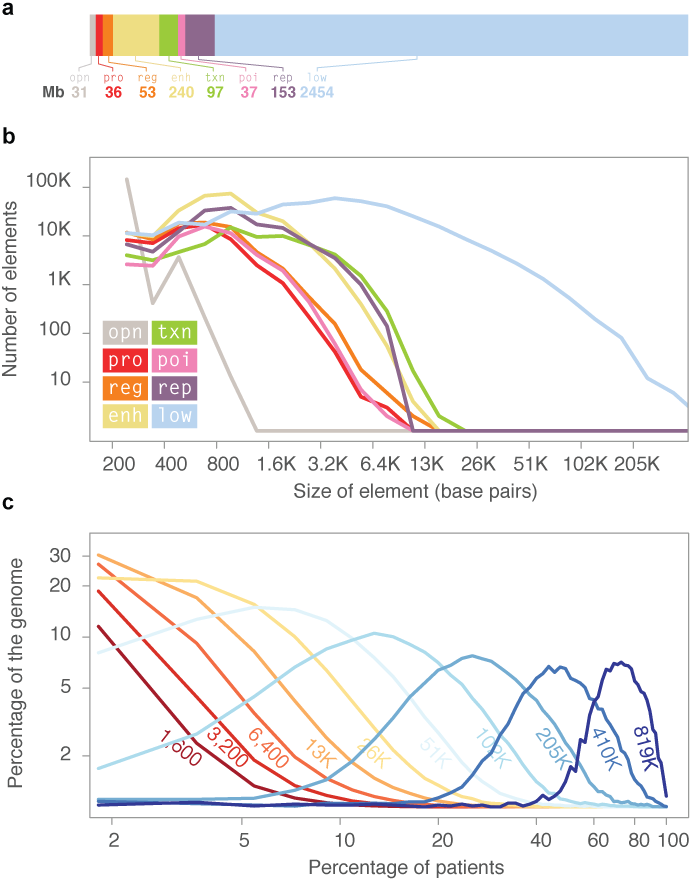
Recurrence at contiguous genomic elements is rare. a. Genome coverage for each of the eight aggregate chromatin states in prostate tissue (RWPE1). b. Number of elements (y-axis) and size of elements (x-axis) for each of the eight chromatin states, showing the distribution of regulatory element size in prostate tissue. The majority of regulatory elements are a few Kb in size. c, Percentage of measurements (y-axis) obtained from sliding a window of variable length (1,6 to 819 Kb) and calculating the percentage of all tumors that harbor at least one mutation in that window (x-axis). Sliding a window of 13 Kb (orange), the size of the largest regulatory elements, yields a distribution of recurrence events across all tumors that rarely exceeds 15% of prostate tumor samples.

**Figure S3.**
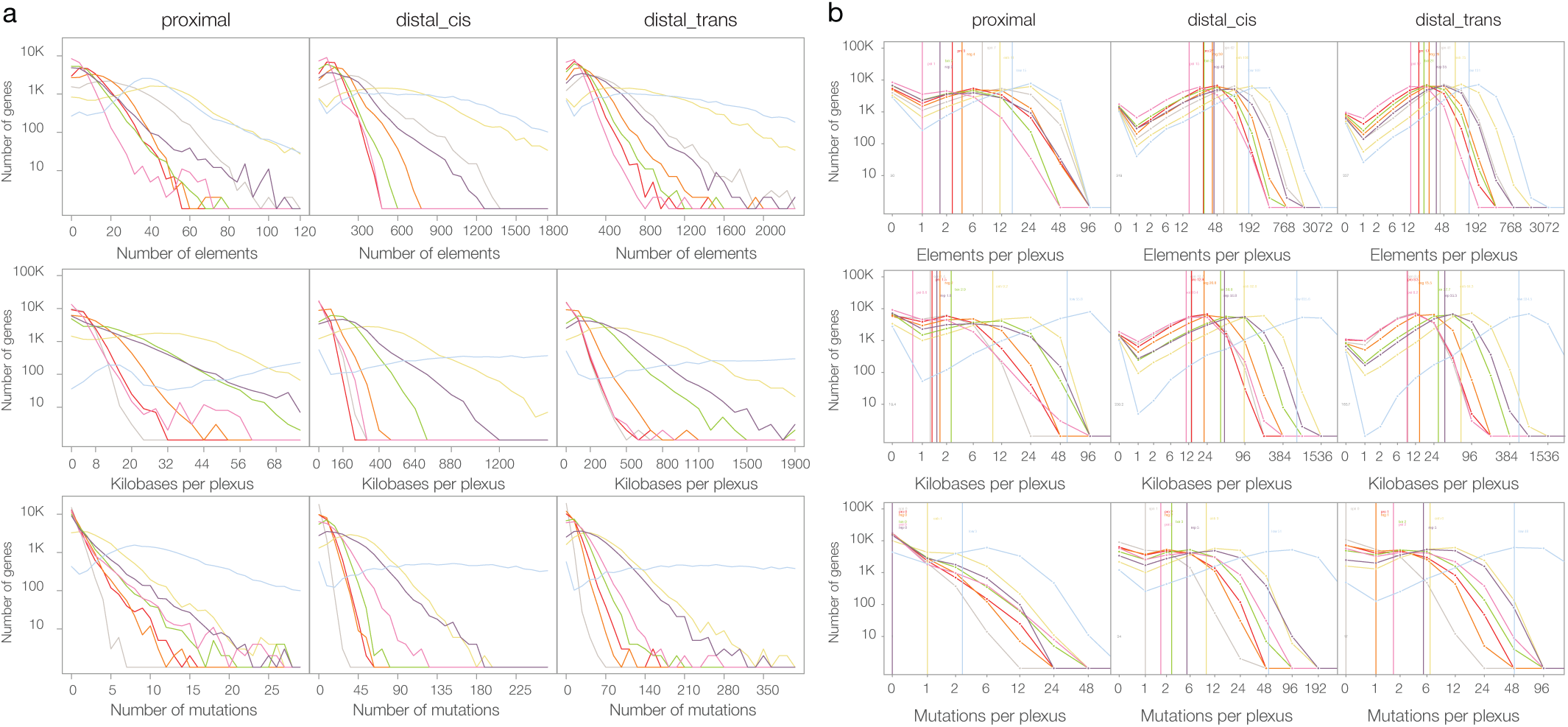
Plexus composition across all genes. a and b. Number of genes (y-axis) and number of elements, nucleotides or mutations for raw (a) or cut (b) plexi (x-axes) for each of the eight aggregate states that are proximal (left), distal intra-chromosomal (*cis*, middle) or distal inter-chromosomal (*trans*, right) with respect to the gene body.

**Figure S4.**
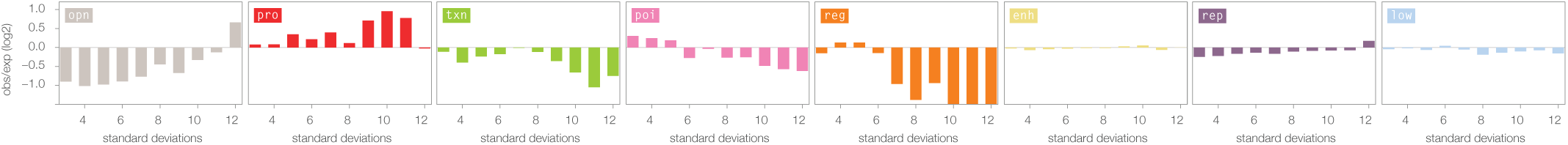
Enrichment scores for mutations in the distal loci of plexi containing dysregulated genes (cut plexi: high-confidence, intra-chromosomal interactions) but excluding gene instances with copy number alterations. Histograms show the log_2_ ratio of observed over expected mutation counts (y-axis) at increasing levels of dysregulation (x-axis).

**Figure S5.**
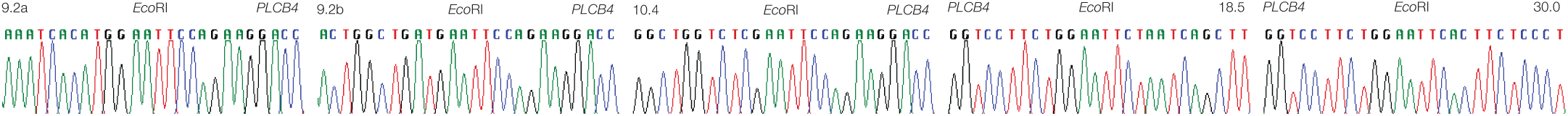
*PLCB4* plexus 3C chromatograms. Sanger sequence chromatograms of 3C ligation products formed between the ***PLBC4*** promoter and mutated genomic loci.

## SUPPLEMENTARY TABLES

**Table S1.**
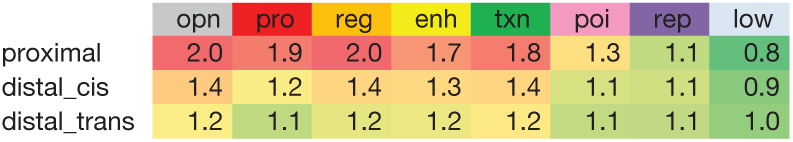
Plexus of protein-coding genes are enriched for regulatory chromatin states. Ratio between the number of tiles observed on average for each plexus and the number of tiles expected based on genomic proportions in prostate cancer.

**Table S2.**
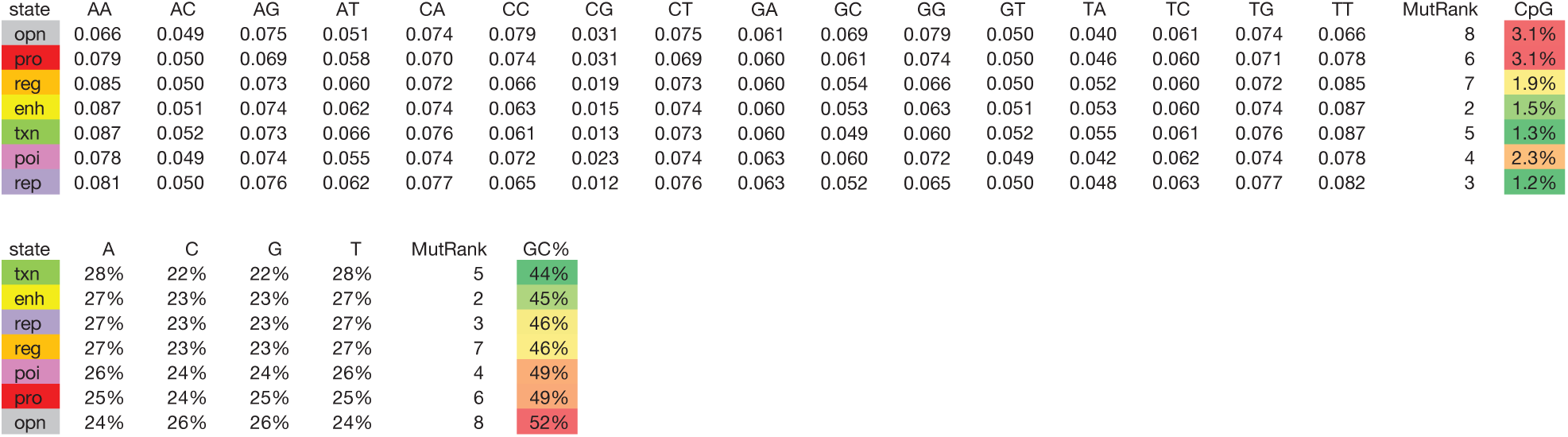
Nucleotide and dinucleotide composition of chromatin states in prostate tissue.

**Table S3.**
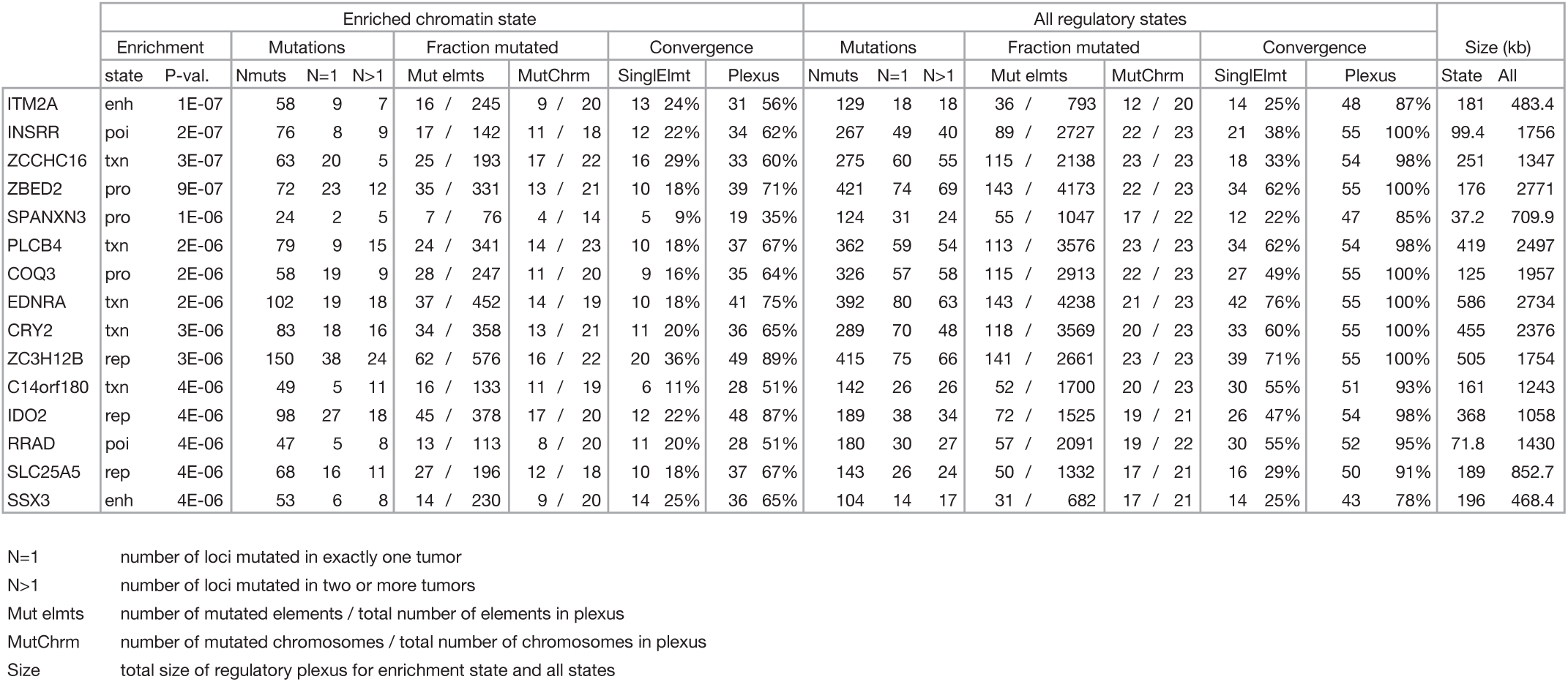
Significant results for the plexus recurrence test run on all GENCODE protein-coding genes.

**Table S4.**
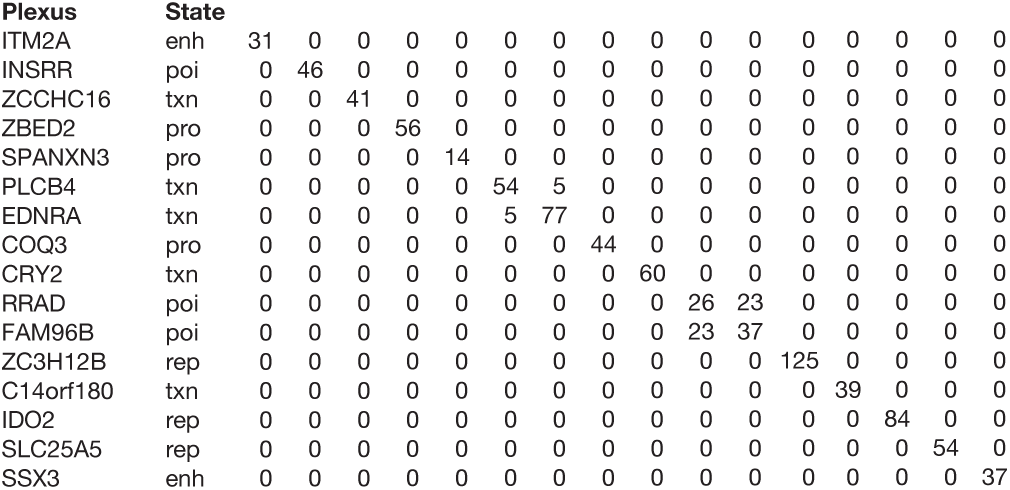
Plexus intersections. Number of shared tiles between all pairs of plexi identified by the plexus recurrence test. Only the portions of the plexus marked by the significantly mutated chromatin state are intersected.

**Table S5.**
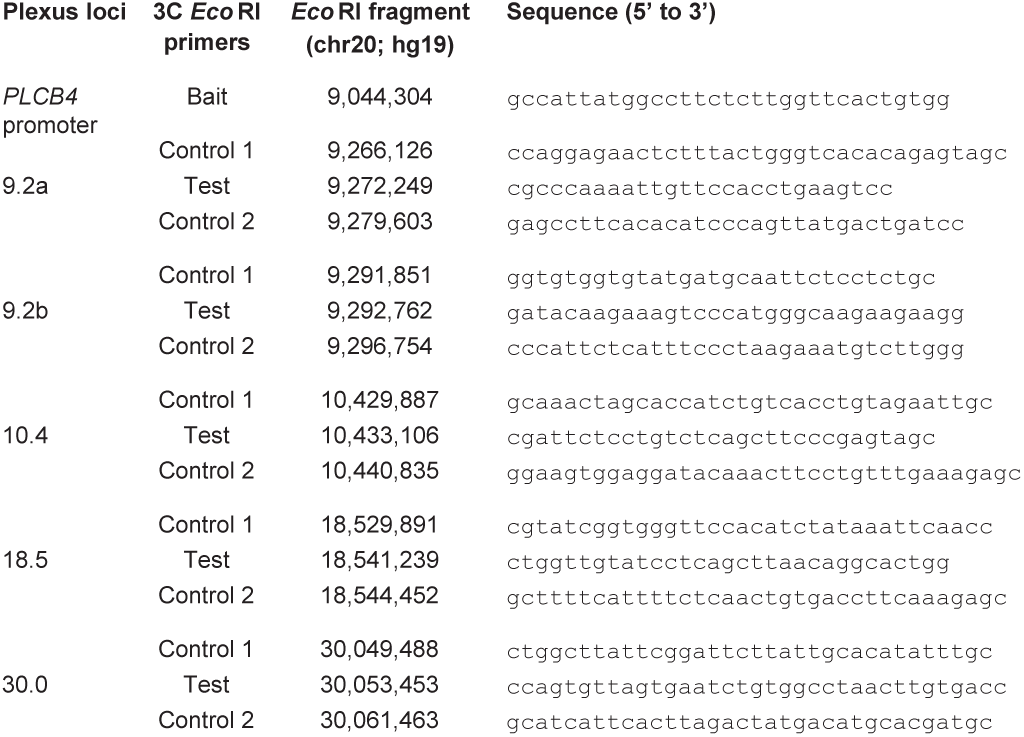
*PLCB4* oligonucleotides. Nucleotide sequence of primers used in chromatin conformation capture (3C) experiments validating the *PLCB4* plexus interactions.

## SUPPLEMENTARY DATA

**Data S1** | Number of interactions in the plexi of protein coding genes. Table header: Internal gene ID, GENCODE gene symbol, Gene class, Number of unique anchors at TSS, Number of proximal interactions, Number of distal cis interactions, Number of distal trans interactions.

**Data S2** | Number of tiles, elements and mutations in the plexi of proteincoding genes. Table header: Internal gene ID, GENCODE gene symbol, Stringency class, Annotation class, Distance class, opn, pro, reg, enh, txn, poi, rep, low.

**Data S3** | Dysregulated-unchanged gene-instance pairs. Table header: Internal gene ID, Gene normal core median, Gene normal core SD, Dysregulated index, Dysregulated patient ID, Dysregulated core score, Unchanged index, Unchanged patient ID, Unchanged core score.

**Data S4** | Plexus recurrence test results for all GENCODE protein-coding genes. Table header: Most significant p-value, Internal gene ID, opn, pro, reg, enh, txn, poi, rep, low, exn, Ensembl gene ID, Gene symbol.

**Data S5** | ***PLCB4*** 3C plexus eQTL results in normal prostate. Table header: Chromosome, Start, End, Gene isoform, Beta, T-statistic, P-value, FDR.

## PLEXUS FRAMEWORK PRESENTATIONS

Cowper [Sallari] Richard, Sinnott-Armstrong Nicholas A, Lupien Mathieu, Kellis, Manolis. “Hierarchical models of mutational recurrence and allelic burden in prostate cancer.” (Abstract 3451T) Presented at the 63rd Annual Meeting of The American Society of Human Genetics, October 22-26, 2013, Boston, MA.

http://www.ashg.org/2013meeting/abstracts/fulltext/f130122912.htm

Sallari, Richard C. “Convergence of dispersed regulatory mutations reveals candidate driver genes in prostate cancer.” Presented at the Banff International Research Station for Mathematical Innovation and Discovery, August 6, 2015, DOI: 10.14288/1.0300033 http://www.birs.ca/events/2015/5-day-workshops/15w5142/videos/watch/201508061144-Sallari.html

## REFERENCES

Ahmadiyeh, Nasim, Mark M Pomerantz, Chiara Grisanzio, Paula Herman, Li Jia, Vanessa Almendro, Housheng Hansen He, et al. 2010. “8q24 Prostate, Breast, and Colon Cancer Risk Loci Show Tissue-Specific Long-Range Interaction with MYC.” Proceedings of the National Academy of Sciences 107 (21): 9742–46. DOI:10.1073/pnas.0910668107.

Akhtar-Zaidi, Batool, Richard Cowper-Sallari, Olivia Corradin, Alina Saiakhova, Cynthia F Bartels, Dheepa Balasubramanian, Lois Myeroff, et al. 2012. “Epigenomic Enhancer Profiling Defines a Signature of Colon Cancer.” Science 336 (6082): 736–39. DOI:10.1126/science.1217277.

Asangani, Irfan A, Vijaya L Dommeti, Xiaoju Wang, Rohit Malik, Marcin Cieslik, Rendong Yang, June Escara-Wilke, et al. 2014. “Therapeutic Targeting of BET Bromodomain Proteins in Castration-Resistant Prostate Cancer.” Nature 510 (7504): 278–82. DOI:10.1038/nature13229.

Baca, Sylvan C, Davide Prandi, Michael S Lawrence, Juan Miguel Mosquera, Alessandro Romanel, Yotam Drier, Kyung Park, et al. 2013. “Punctuated Evolution of Prostate Cancer Genomes.” Cell 153 (3): 666–77. DOI:10.1016/j.-cell.2013.03.021.

Barbieri, Christopher E, Sylvan C Baca, Michael S Lawrence, Francesca Demichelis, Mirjam Blattner, Jean-Philippe Theurillat, Thomas A White, et al. 2012. “Exome Sequencing Identifies Recurrent SPOP, FOXA1 and MED12 Mutations in Prostate Cancer.” Nature Genetics 44 (6): 685–89. DOI:10.1038/ng.2279.

Chu, L W, Y Zhu, K Yu, T Zheng, H Yu, Y Zhang, I Sesterhenn, et al. 2007. “Variants in Circadian Genes and Prostate Cancer Risk: a Population-Based Study in China.” Prostate Cancer and Prostatic Diseases 11 (4): 342–48. DOI:10.1038/sj.pcan.4501024.

Cowper-Sallari, Richard, Xiaoyang Zhang, Jason B Wright, Swneke D Bailey, Michael D Cole, Jerome Eeckhoute, Jason H Moore, and Mathieu Lupien. 2012. “Breast Cancer Risk-Associated SNPs Modulate the Affinity of Chromatin for FOXA1 and Alter Gene Expression.” Nature Genetics 44 (11): 1191–98. DOI:10.1038/ng.2416.

D’Arcy, Padraig, Wessen Maruwge, Barry Wolahan, Limin Ma, and Bertha Brodin. 2014. “Oncogenic Functions of the Cancer-Testis Antigen SSX on the Proliferation, Survival, and Signaling Pathways of Cancer Cells.” PloS one 9 (4): e95136. DOI:10.1371/journal.pone.0095136.

ENCODE Consortium. 2012. “An Integrated Encyclopedia of DNA Elements in the Human Genome.” Nature 489 (7414): 57–74. DOI:10.1038/nature11247.

Ernst, Jason, and Manolis Kellis. 2012. “ChromHMM: Automating Chromatin-State Discovery and Characterization.” Nature Methods 9 (3): 215–16. DOI:10.1038/nmeth.1906.

Ernst, Jason, and Manolis Kellis. 2013. “Interplay Between Chromatin State, Regulator Binding, and Regulatory Motifs in Six Human Cell Types.” Genome research 23 (7). 1142–54. DOI:10.1101/gr.144840.112.

Ernst, Jason, and Manolis Kellis. 2015. “Large-Scale Imputation of Epigenomic Datasets for Systematic Annotation of Diverse Human Tissues.” Nature Biotechnology 33 (4): 364–76. DOI:10.1038/nbt.3157.

Fuchsberger, Christian, Jason Flannick, Tanya M Teslovich, Anubha Mahajan, Vineeta Agarwala, Kyle J Gaulton, Clement Ma, et al. 2016. “The Genetic Architecture of Type 2 Diabetes.” Nature 536 (7614): 41–47. DOI:10.1038/nature18642.

Griffon, Aurélien, Quentin Barbier, Jordi Dalino, Jacques van Helden, Salvatore Spicuglia, and Benoit Ballester. 2015. “Integrative Analysis of Public ChIP-Seq Experiments Reveals a Complex Multi-Cell Regulatory Landscape.” Nucleic Acids Research 43 (4): e27–e27. DOI:10.1093/nar/gku1280.

GTEx Consortium. 2015. “Human Genomics. the Genotype-Tissue Expression (GTEx) Pilot Analysis: Multitissue Gene Regulation in Humans.” Science 348 (6235): 648–60. DOI:10.1126/science.1262110.

Hua, Xing, Haiming Xu, Yaning Yang, Jun Zhu, Pengyuan Liu, and Yan Lu. 2013. “DrGaP: a Powerful Tool for Identifying Driver Genes and Pathways in Cancer Sequencing Studies.” The American Journal of Human Genetics 93 (3): 439–51. DOI:10.1016/j.ajhg.2013.07.003.

Huang, Franklin W, Eran Hodis, Mary Jue Xu, Gregory V Kryukov, Lynda Chin, and Levi A Garraway. 2013. “Highly Recurrent TERT Promoter Mutations in Human Melanoma.” Science 339 (6122): 957–59. DOI:10.1126/-science.1229259.

Khurana, Ekta, Yao Fu, Vincenza Colonna, Xinmeng Jasmine Mu, Hyun Min Kang, Tuuli Lappalainen, Andrea Sboner, et al. 2013. “Integrative Annotation of Variants From 1092 Humans: Application to Cancer Genomics.” Science 342 (6154): 1235587. DOI:10.1126/science.1235587.

Lawrence, Michael S, Petar Stojanov, Craig H Mermel, James T Robinson, Levi A Garraway, Todd R Golub, Matthew Meyerson, Stacey B Gabriel, Eric S Lander, and Gad Getz. 2014. “Discovery and Saturation Analysis of Cancer Genes Across 21 Tumour Types.” Nature 505 (7484): 495–501. DOI:10.1038/-nature12912.

Lawrence, Michael S, Petar Stojanov, Paz Polak, Gregory V Kryukov, Kristian Cibulskis, Andrey Sivachenko, Scott L Carter, et al. 2013. “Mutational Heterogeneity in Cancer and the Search for New Cancer-Associated Genes.” Nature 499 (7457): 214–18. DOI:10.1038/nature12213.

Liu, Lin, Subhajyoti De, and Franziska Michor. 2013. “DNA Replication Timing and Higher-Order Nuclear Organization Determine Single-Nucleotide Substitution Patterns in Cancer Genomes.” Nature Communications 4: 1502. DOI:10.1038/ncomms2502.

Lupien, Mathieu, Jerome Eeckhoute, Clifford A Meyer, Qianben Wang, Yong Zhang, Wei Li, Jason S Carroll, X Shirley Liu, and Myles Brown. 2008. “FoxA1 Translates Epigenetic Signatures Into Enhancer-Driven Lineage-Specific Transcription.” Cell 132 (6): 958–70. DOI:10.1016/j.cell.2008.01.018.

Mack, S C, H Witt, R M Piro, L Gu, S Zuyderduyn, A M Stütz, X Wang, et al. 2014. “Epigenomic Alterations Define Lethal CIMP-Positive Ependymomas of Infancy.” Nature 506 (7489): 445–50. DOI:10.1038/nature13108.

Magnani, Luca, Alexander Stoeck, Xiaoyang Zhang, András Lánczky, Anne C Mirabella, Tian-Li Wang, Balázs Gyorffy, and Mathieu Lupien. 2013. “Genome-Wide Reprogramming of the Chromatin Landscape Underlies Endocrine Therapy Resistance in Breast Cancer.” 110 (16). Proceedings of the National Academy of Sciences 110 (16): E1490–99. DOI:10.1073/pnas.1219992110.

Metz, Richard, Courtney Smith, James B DuHadaway, Phillip Chandler, Babak Baban, Lauren M F Merlo, Elizabeth Pigott, et al. 2014. “IDO2 Is Critical for IDO1-Mediated T-Cell Regulation and Exerts a Non-Redundant Function in Inflammation.” International Immunology 26 (7): 357–67. DOI:10.1093/intimm/dxt073.

Nik-Zainal, Serena, Helen Davies, Johan Staaf, Manasa Ramakrishna, Dominik Glodzik, Xueqing Zou, Inigo Martincorena, et al. 2016. “Landscape of Somatic Mutations in 560 Breast Cancer Whole-Genome Sequences.” Nature 534 (7605): 47–54. DOI:10.1038/nature17676.

Park, Donghyun, Sukchan Lee, Kisun Jun, Yeon-Mi Hong, Do Young Kim, Yang In Kim, and Hee-Sup Shin. 2003. “Translation of Clock Rhythmicity Into Neural Firing in Suprachiasmatic Nucleus Requires mGluR-PLCbeta4 Signaling.” Nature Neuroscience 6 (4): 337–38. DOI:10.1038/nn1033.

Patel, B, Y Kang, K Cui, M Litt, M S J Riberio, C Deng, T Salz, et al. 2014. “Aberrant TAL1 Activation Is Mediated by an Interchromosomal Interaction in Human T-Cell Acute Lymphoblastic Leukemia.” Leukemia 28 (2): 349–61. DOI:10.1038/leu.2013.158.

Polak, Paz, Michael S Lawrence, Eric Haugen, Nina Stoletzki, Petar Stojanov, Robert E Thurman, Levi A Garraway, et al. 2014. “Reduced Local Mutation Density in Regulatory DNA of Cancer Genomes Is Linked to DNA Repair.” Nature Biotechnology 32 (1): 71–75. DOI:10.1038/nbt.2778.

Polak, Paz, Robert Querfurth, and Peter F Arndt. 2010. “The Evolution of Transcription-Associated Biases of Mutations Across Vertebrates.” BMC Evolutionary Biology 10 (1): 187. DOI:10.1186/1471-2148-10-187.

Pomerantz, Mark M, Nasim Ahmadiyeh, Li Jia, Paula Herman, Michael P Verzi, Harshavardhan Doddapaneni, Christine A Beckwith, et al. 2009. “The 8q24 Cancer Risk Variant Rs6983267 Shows Long-Range Interaction with MYC in Colorectal Cancer.” Nature Genetics 41 (8): 882–84. DOI:10.1038/ng.403.

Reynet, C, and C R Kahn. 1993. “Rad: a Member of the Ras Family Overexpressed in Muscle of Type II Diabetic Humans.” Science 262 (5138): 1441–44. DOI:10.1126/science.8248782.

Rickman, David S, T David Soong, Benjamin Moss, Juan Miguel Mosquera, Jan Dlabal, Stéphane Terry, Theresa Y MacDonald, et al. 2012. “Oncogene-Mediated Alterations in Chromatin Conformation.” Proceedings of the National Academy of Sciences 109 (23): 9083–9088. DOI:10.1073/pnas.1112570109.

Rieder, Mark J, Glenn E Green, Sarah S Park, Brendan D Stamper, Christopher T Gordon, Jason M Johnson, Christopher M Cunniff, et al. 2012. “A Human Homeotic Transformation Resulting From Mutations in PLCB4 and GNAI3 Causes Auriculocondylar Syndrome.” American Journal of Human Genetics 90 (5): 907–14. DOI:10.1016/j.ajhg.2012.04.002.

Roadmap Epigenomics Consortium, Anshul Kundaje, Wouter Meuleman, Jason Ernst, Misha Bilenky, Angela Yen, Alireza Heravi-Moussavi, et al. 2015. “Integrative Analysis of 111 Reference Human Epigenomes.” Nature 518 (7539): 317–30. DOI:10.1038/nature14248.

Romanuik, Tammy L, Gang Wang, Olena Morozova, Allen Delaney, Marco A Marra, and Marianne D Sadar. 2010. “LNCaP Atlas: Gene Expression Associated with in Vivo Progression to Castration-Recurrent Prostate Cancer.” BMC Medical Genomics 3 (1): 43. DOI:10.1186/1755-8794-3-43.

Sankpal, Umesh T, Steven Goodison, Maen Abdelrahim, and Riyaz Basha. 2011. “Targeting Sp1 Transcription Factors in Prostate Cancer Therapy.” Medicinal Chemistry 7 (5): 518–25.

Sheridan, Cormac. 2015. “IDO Inhibitors Move Center Stage in Immuno-Oncology.” Nature Biotechnology 33: 321–322. DOI:10.1038/nbt0415-321.

Stamatoyannopoulos, John A, Ivan Adzhubei, Robert E Thurman, Gregory V Kryukov, Sergei M Mirkin, and Shamil R Sunyaev. 2009. “Human Mutation Rate Associated with DNA Replication Timing.” Nature Genetics 41 (4): 393–95. DOI:10.1038/ng.363.

Tomlins, Scott A, Daniel R Rhodes, Sven Perner, Saravana M Dhanasekaran, Rohit Mehra, Xiao-Wei Sun, Sooryanarayana Varambally, et al. 2005. “Recurrent Fusion of TMPRSS2 and ETS Transcription Factor Genes in Prostate Cancer.” Science 310 (5748): 644–48. DOI:10.1126/-science.1117679.

Tucci, Paola, Massimiliano Agostini, Francesca Grespi, Elke K Markert, Alessandro Terrinoni, Karen H Vousden, Patricia A J Muller, et al. 2012. “Loss of P63 and Its microRNA-205 Target Results in Enhanced Cell Migration and Metastasis in Prostate Cancer.” Proceedings of the National Academy of Sciences 109 (38): 15312–17. DOI:10.1073/pnas.1110977109.

Zhang, Xiaoyang, Richard Cowper-Sal·lari, Swneke D Bailey, Jason H Moore, and Mathieu Lupien. 2012. “Integrative Functional Genomics Identifies an Enhancer Looping to the SOX9 Gene Disrupted by the 17q24.3 Prostate Cancer Risk Locus.” Genome Research 22 (8): 1437–46. DOI:10.1101/gr.135665.111.

Zhu, Yong, Richard G Stevens, Aaron E Hoffman, Liesel M FitzGerald, Erika M Kwon, Elaine A Ostrander, Scott Davis, Tongzhang Zheng, and Janet L Stanford. 2009. “Testing the Circadian Gene Hypothesis in Prostate Cancer: a Population-Based Case-Control Study.” Cancer Research 69 (24): 9315–22. DOI:10.1158/0008-5472.CAN-09-0648.

## REFERENCES

Baca, Sylvan C, Davide Prandi, Michael S Lawrence, Juan Miguel Mosquera, Alessandro Romanel, Yotam Drier, Kyung Park, et al. 2013. “Punctuated Evolution of Prostate Cancer Genomes.” Cell 153 (3): 666–77. doi:10.1016/j.cell.2013.03.021.

Bailey, Swneke D, Xiaoyang Zhang, Kinjal Desai, Malika Aid, Olivia Corradin, Richard Cowper-Sal-lari, Batool Akhtar-Zaidi, Peter C Scacheri, Benjamin Haibe-Kains, and Mathieu Lupien. 2015. “ZNF143 Provides Sequence Specificity to Secure Chromatin Interactions at Gene Promoters.” Nature Communications 2: 6186. doi:10.1038/ncomms7186.

Cowper-Sal-lari, Richard, Xiaoyang Zhang, Jason B Wright, Swneke D Bailey, Michael D Cole, Jerome Eeckhoute, Jason H Moore, and Mathieu Lupien. 2012. “Breast Cancer Risk-Associated SNPs Modulate the Affinity of Chromatin for FOXA1 and Alter Gene Expression.” Nature Genetics 44 (11). Nature Research: 1191–98. doi:10.1038/ng.2416.

Ernst, Jason, and Manolis Kellis. 2012. “ChromHMM: Automating Chromatin-State Discovery and Characterization.” Nature Methods 9 (3): 215–16. doi:10.1038/nmeth.1906.

Ghoussaini, Maya, and Stacey L. Edwards. 2014. “Evidence that breast cancer risk at the 2q35 locus is mediated through IGFBP5 regulation.” Nature communications 4: 4999. doi:10.1038/ncomms5999.

GTEx Consortium. 2015. “Human Genomics. the Genotype-Tissue Expression (GTEx) Pilot Analysis: Multitissue Gene Regulation in Humans.” Science 348 (6235): 648–60. doi:10.1126/science.1262110.

Lawrence, Michael S, Petar Stojanov, Paz Polak, Gregory V Kryukov, Kristian Cibulskis, Andrey Sivachenko, Scott L Carter, et al. 2013. “Mutational Heterogeneity in Cancer and the Search for New Cancer-Associated Genes.” Nature 499 (7457): 214–18. doi:10.1038/nature12213.

Ramskold, Daniel, Eric T. Wang, Christopher B. Burge, and Rickard Sandberg. 2009. “An abundance of ubiquitously expressed genes revealed by tissue transcriptome sequence data.” PLoS computational biology 5, no. 12: e1000598. doi:10.1371/journal.pcbi.1000598.

Rickman, David S, T David Soong, Benjamin Moss, Juan Miguel Mosquera, Jan Dlabal, Stéphane Terry, Theresa Y MacDonald, et al. 2012. “Oncogene-Mediated Alterations in Chromatin Conformation.” PNAS 109 (23): 9083–88. doi:10.1073/pnas.1112570109.

Shabalin, Andrey A. 2012. “Matrix eQTL: Ultra Fast eQTL Analysis via Large Matrix Operations.” Bioinformatics 28 (10): 1353–58. doi:10.1093/bioinformatics/bts163.

Sharma, Naomi L, Charlie E Massie, Falk Butter, Matthias Mann, Helene Bon, Antonio Ramos-Montoya, Suraj Menon, et al. 2014. “The ETS Family Member GABPα Modulates Androgen Receptor Signalling and Mediates an Aggressive Phenotype in Prostate Cancer.” Nucleic Acids Research 42 (10): 6256–69. doi:10.1093/nar/gku281.

